# Three-Dimensional Mitochondria Reconstructions of Murine Cardiac Muscle Changes in Size Across Aging

**DOI:** 10.1101/2022.04.23.489291

**Authors:** Zer Vue, Kit Neikirk, Larry Vang, Edgar Garza-Lopez, Trace A. Christensen, Jianqiang Shao, Jacob Lam, Heather K. Beasley, Andrea G. Marshall, Amber Crabtree, B Josephs, Benjamin Rodriguez, Benjamin Kirk, Serif Bacevac, Taylor Barongan, Bryanna Shao, Dominique C. Stephens, Kinuthia Kabugi, Ho-Jin Koh, Alice Koh, Chantell S. Evans, Brittany Taylor, Anilkumar K. Reddy, Tyne Miller-Flemming, Ky’Era V. Actkins, Elma Zaganjor, Nastaran Daneshgar, Sandra A. Murray, Bret C. Mobley, Steven Damo, Jennifer A. Gaddy, Blake Riggs, Celestine Wanjalla, Annet Kirabo, Melanie McReynolds, Jose A. Gomez, Mark A. Phillips, Vernat Exil, Dao-Fu Dai, Antentor Hinton

**Affiliations:** Department of Molecular Physiology and Biophysics, Vanderbilt University, Nashville, TN, 37232, USA; Department of Internal Medicine, University of Iowa, Iowa City, IA, 52242, USA; Microscopy and Cell Analysis Core Facility, Mayo Clinic, Rochester, MN, 55905, USA; Central Microscopy Research Facility, University of Iowa, Iowa City, IA, 52242, USA; Department of Life and Physical Sciences, Fisk University, Nashville, TN, 37208, USA; Department of Biological Sciences, Tennessee State University, Nashville, TN 37209, USA; Department of Cell Biology, Duke University School of Medicine, Durham, NC, 27709, USA; J. Crayton Pruitt Family Department of Biomedical Engineering, University of Florida, Gainesville, FL, 32612, USA; Department of Medicine, Baylor College of Medicine, Houston, TX, 77002, USA; Division of Genetic Medicine, Department of Medicine, Vanderbilt University Medical Center, Nashville, TN, USA; Department of Pathology, Carver College of Medicine, University of Iowa, Iowa City, IA, 52243, USA; Department of Cell Biology, School of Medicine, University of Pittsburgh, Pittsburgh, PA, 15261, USA; Department of Pathology, Microbiology and Immunology, Vanderbilt University Medical Center, Nashville, TN, 37233 USA; Department of Medicine, Vanderbilt University Medical Center, Nashville, TN 37232, USA; Tennessee Valley Healthcare Systems, U.S. Department of Veterans Affairs, Nashville, TN 37212, USA; Department of Biology at San Francisco State University, San Francisco, CA 94133, USA; Department of Medicine / Clinical Pharmacology Division. Vanderbilt University Medical Center, Nashville, TN 37232; Department of Biochemistry and Molecular Biology, Pennsylvania State University, State College, PA 16801; Department of Medicine, Vanderbilt University Medical Center, Nashville, TN, 37233, USA; Department of Integrative Biology, Oregon State University, Corvallis, OR, 97332, USA; Department of Pediatrics, Div. of Cardiology, St. Louis University School of Medicine, St. Louis, MO, 63105, USA; Department of Pediatrics, Carver College of Medicine, University of Iowa, Iowa City, IA 52243, USA; Department of Pathology, University of John Hopkins, Baltimore, MD, USA

**Keywords:** MICOS, aging, mitochondria, 3D morphometry, mitochondrial disease, mitochondrion, reconstruction, reticulum, serial block-face SEM, cardiac muscle

## Abstract

With sparse treatment options, cardiac disease remains a significant cause of death among humans. As a person ages, mitochondria break down and the heart becomes less efficient. Heart failure is linked to many mitochondria-associated processes, including endoplasmic reticulum stress, mitochondrial bioenergetics, insulin signaling, autophagy, and oxidative stress. The roles of key mitochondrial complexes that dictate the ultrastructure, such as the mitochondrial contact site and cristae organizing system (MICOS), in aging cardiac muscle are poorly understood. To better understand the cause of age-related alteration in mitochondrial structure in cardiac muscle, we used transmission electron microscopy (TEM) and serial block facing-scanning electron microscopy (SBF-SEM) to quantitatively analyze the 3D networks in cardiac muscle samples of male mice at aging intervals of 3 months, 1 year, and 2 years. Here, we present the loss of cristae morphology, the inner folds of the mitochondria, across age. In conjunction with this, the 3D volume of mitochondria decreased. These findings mimicked observed phenotypes in murine cardiac fibroblasts with CRISPR/Cas9 knockout of *Mitofilin, Chchd3, Chchd6* (some members of the MICOS complex), and *Opa1*, which showed poorer oxidative consumption rate and mitochondria with decreased mitochondrial length and volume. In combination, these data show the need to explore if loss of the MICOS complex in the heart may be involved in age-associated mitochondrial and cristae structural changes.

## INTRODUCTION

The mitochondrion is an organelle that critically carries out oxidative phosphorylation in the cell and is implicated in many diseases, including Alzheimer’s disease, diabetes, and heart failure (1–3). Given the dependence on mitochondria to meet energetic demands, mitochondrial dysfunction and deterioration are associated with cell death, including autophagy and apoptosis (4–6). In addition to producing adenosine triphosphate (ATP), mitochondria also regulate cell signaling during various cellular processes, including calcium homeostasis, apoptosis, and immunity (5, 7–9). Interestingly, research also suggests that mitochondria play a significant role in aging (10, 11). Although the functions of mitochondria are well understood, the relationship between mitochondrial dysfunction and age-associated diseases requires further investigation.

Mitochondrial morphology is associated with mitochondrial bioenergetic capacity Mitochondrial dynamics, mainly fusion and fission, provide a means for mitochondria to readily adapt to the specific energy demands of the cell. Fusion and fission are regulated by optic atrophy 1 (OPA1) and dynamin-related protein-1 (DRP1), respectively (4, 12–14). While the loss of DRP1 results in an abundance of overly elongated and fused mitochondria (5), the loss of OPA1 gives rise to fragmented mitochondria (12, 15). Mitochondria may go through several forms in coordinated cycles of fusion and fission. If OPA1 and mitofusions become upregulated, fusion rates increase producing long, tubular mitochondria with more complex network formation (16). Under normal physiological conditions, tubular mitochondria function well. However, under conditions of stress, mitochondria have been shown to alter their formation into donut, blob, or fragmented shapes, the latter of which is caused by a high fission-fusion ratio (16–18). Importantly, the evaluation of mitochondrial structure is vital as deviations from typical tubular mitochondrial shape indicate lower ATP production and increased susceptibility to autophagosome degradation (19). Moreover, the loss of several mitochondrial genes has been implicated in cristae structure alterations (13, 20).

Cristae, the folds of the mitochondria inner membrane, house electron transfer chain complexes responsible for establishing the proton motive force, which ultimately produces ATP (21). The morphological arrangement of cristae ultimately determines the energetic capacity of mitochondria and has been implicated in playing a role in other functions including apoptosis and homeostasis (21). The mitochondrial contact site and cristae organizing system (MICOS) maintains the morphology of cristae (22). Similar to OPA1, the MICOS complex governs the dynamics of cristae structure (20, 22–24). The MICOS complex is also involved in the maintenance of cristae junctions, which are sub-compartments for metabolites (25). Given the significance of cristae junctions, cristae may also be involved in other processes such as regulating calcium homeostasis and molecular signaling. Considering the significance of mitochondrial and cristae morphology in normal function, dysfunction of the MICOS complex could negatively impact health (24). However, the role of the MICOS complex in mitochondrial health during cardiac aging has yet to be defined. Given the understanding that mitochondria are implicated in the aging process (11), we hypothesized that changes in the MICOS complex, which is critical to maintaining cristae morphology (26, 27), may lead to age-associated changes in mitochondria.

Specifically, we sought to understand the role of the MICOS complex in cardiac muscle mitochondria as it may provide novel insights into maintaining normal mitochondrial health in the heart. This study combines an *in vivo* and *in vitro* approach that looks at cardiac muscle in mice, cardiac fibroblasts, and human induced pluripotent stem cell-derived cardiomyocytes (iPSC-CMs). Previously, mitochondrial-focused aging studies have focused on cardiomyocytes (28); however, these studies overlooked the MICOS complex in relation to the aging of the cardiac fibroblasts. Cardiac health is linked to insulin resistance and diabetes (1, 29). Heart failure is also associated with other mitochondrial processes (6, 30–32) such as endoplasmic reticulum stress, mitochondrial bioenergetics, insulin signaling, autophagy, and oxidative stress (33). Mitochondria have specialized types and structures associated with their roles in glycolysis and oxidative metabolism in cardiac fibroblasts (17, 34–36). While young mitochondria show extensive specialization, it is unclear whether this is also true for aged cardiac murine mitochondria, and there is little information quantitively analyzing how aging affects cristae and 3D mitochondrial structure across aging in cardiac tissue. In cardiac muscle, research has shown that across aging, mitochondria in between myofibrils, called intermyofibrillar mitochondria, have the most significant decrease in mitochondrial oxidative respiration function (37). However, the full extent of causes of these changes in the oxidative phosphorylation of intermyofibrillar mitochondria have yet to be elucidated. Given that heart failure risk increases with aging (38), it is vital to consider the role of mitochondrial dysfunction in the loss of heart function (33).

To better understand the relationship between cardiac mitochondrial structure and the aging process, we used transmission electron microscopy (TEM) and fluorescence for 2D micrographs to observe mitochondria and cristae. Different muscle types have distinct mitochondrial function (17, 34). However, it is unclear how these unique functioning differences change over age. To further understand how aging affected mitochondrial morphology, we looked at cristae morphology at three-time points; 3-month, 1-year, and 2-year, which represent “young,” “middle-aged,” and “old,” mice, respectively. We expanded this to include serial block-face scanning electron microscopy (SBF-SEM) with manual contour tracing to reconstruct 3D mitochondrial morphology and structure. We then quantitatively analyzed the 3D networks in mouse cardiac muscle samples at different age intervals. Notably, 3D reconstruction is an important avenue as the 3D morphology of mitochondria has been linked to their functional capacity (17). Using SBF-SEM, we were able to look at single organelles to compare the shape, morphology, count, complexity, and branching across different ages. Furthermore, we compared the aging effects to the loss of the MICOS complex in relation to mitochondrial morphology alterations. Specifically, we used CRISPR/Cas9 on cardiac fibroblasts to knockout (KO) three genes of the MICOS complex: *Chchd3* (Mic19), *Chchd6* (Mic25), and *Mitofilin* (Mic60), as well as *Opa1*, as a positive control, to see how the loss of the MICOS complex affected mitochondrial size, morphology, and oxygen consumption rate. Finally, we sought to understand the role of the MICOS complex, specifically *Chchd6*, in modulating the reactive oxygen species response in iPSC-CMs.

## RESULTS

### Ultrastructural Changes in Mitochondria and Cristae with Age

It is commonly understood that heart function declines with age in human and murine models. However, these changes and their association to mitochondrial morphological changes are unclear. To investigate the cardiac function of aged mice cohorts, prior to TEM analysis, we collected and analyzed echometric data. We observed changes with age that included increased cardiac output, stroke volume, and LV thickness, without significant changes in normalized heart mass, ejection fraction, or heart rate (Figure S1).

As expected, mice increased in body weight across aging (Fig S2A). Specifically, the fat mass increased significantly faster than lean mass, especially in the 2-year sample (Fig S2B-C). In accordance with this, heart mass also significantly increased with age (Fig S2D). After a 6-hour fast, a glucose tolerance test (GTT) showed that aged mice had higher plasma glucose levels post-GTT, indicating impaired glucose tolerance (Fig S2E-F). These results are consistent with aging mice having larger fat depots and abnormal glucose status in comparison to younger mice. Critically, on average, even with decreased glucose tolerance, aged mice did not reach blood glucose levels that are indicative of diabetes. Therefore, this cohort served as a strong representation of age-independent from pathology in this study. To further understand how mitochondrial affected these dynamics, TEM was utilized to understand aging mitochondrial and cristae morphology.

We examined mitochondria and cristae morphology in cardiac muscle at 3-month, one-year, and 2-year-old male mice. Different cell types can be studied in cardiac muscle (36) and cardiac muscle can give information about potential causes of loss of optimal heart function with age (38). TEM is a powerful tool for studying cristae in mitochondria as it gives very high-quality micrographs (39). Young male mice (n=3, per age group) showed very clear mitochondria with electron-dense membranes, whereas the aged mice had fewer mitochondria and cristae (Figure 1A-A’’). The number of mitochondria more than quadrupled between the 3-month to 1-year aged samples before slightly leveling out between 1-year to 2-year (Figure 1B). Although mitochondrial abundance increased, the average mitochondrial area was significantly smaller in samples older than 3-month; the decrease is smaller between the 1-year and 2-year samples (Figure 1C). This shows a compensating effect as mitochondrial area per fiber area does not show significant changes across the total aging process (Figure 1D). The mitochondria also became slightly more circular (Figure 1E). To measure ultrastructural details of mitochondria, cristae numbers in each mitochondrion were evaluated and consistently showed that the number of cristae decreased with age (Figure 1F). Finally, the cristae score was used to assess the cristae quantity and morphology as it is often used to holistically evaluate cristae (39, 40). A cristae score of 0 represents no clearly defined cristae, 1 represents that greater than 50% of mitochondrial area is devoid of cristae, a cristae score of 2 represents that less than 75% of the mitochondria area has cristae, and the maximum cristae score of 4 represents overall typically defined cristae with normal architecture. There was a large drop in the quality of cristae in cardiac muscle with age. The young 3-month sample had mostly regular and slightly irregular cristae, with cristae scores of about 3.3. The aged samples both had cristae scores around 2, which suggests there are many areas lacking cristae or having irregular cristae (Figure 1G). Thus, we found that cristae count and quality both decreased across the aging process. These important findings that indicate a loss of cristae folds with aging caused us to wonder if cristae were differentially affected in a female model (Figure 1H-H’’).

**Figure 1:**
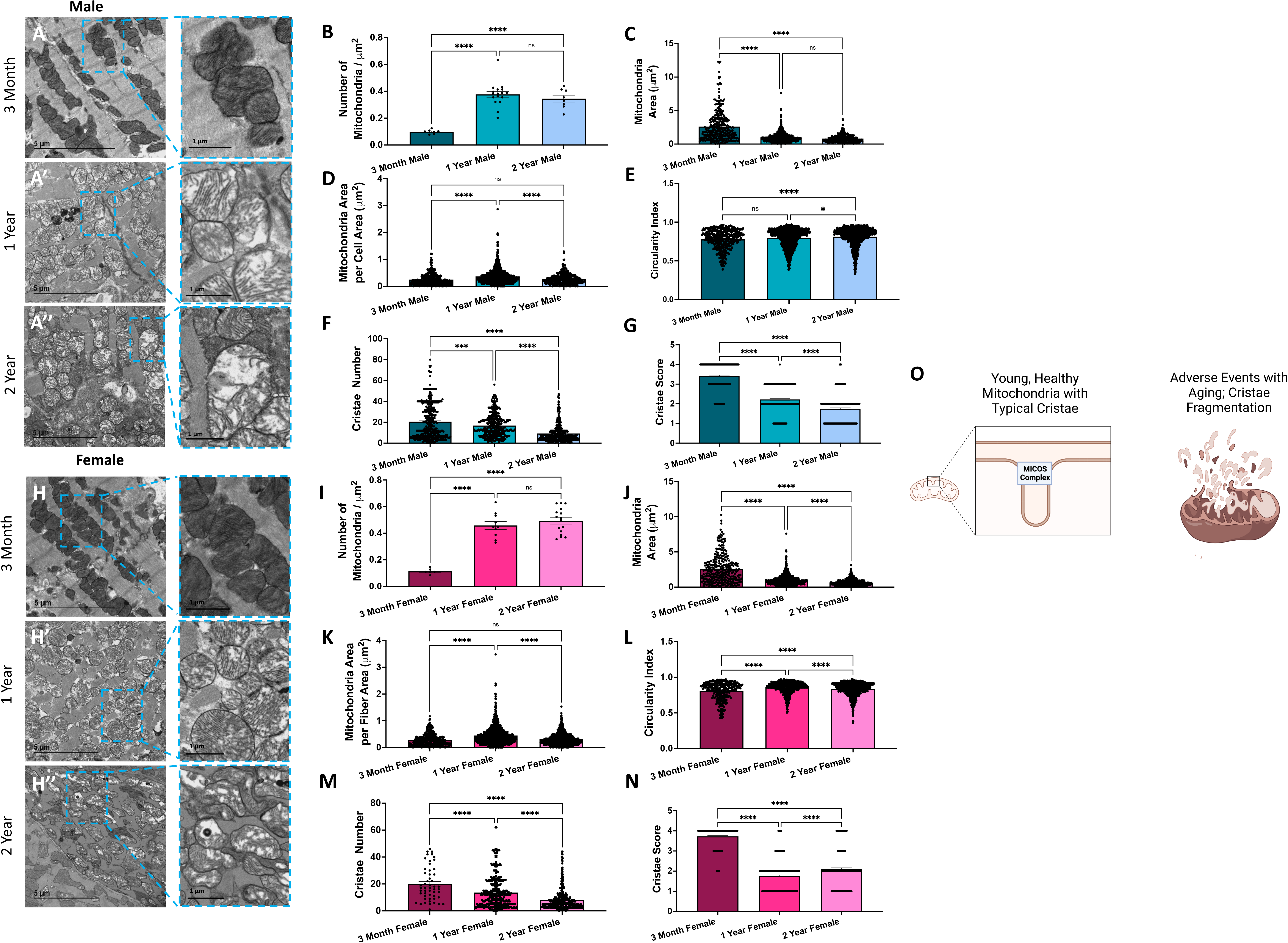
Changes in cardiac muscle mitochondria and cristae across aging revealed in TEM and heart data. Representative transmission electron micrographs for cardiac muscle at (**A**) 3-months, (**A’**) 1-year (**A’’**) and 2-years in male mice. Blue boxes show cristae magnified to enhance the details of cristae. Quantification of key mitochondrial characteristics included (**B**) the number of mitochondria normalized to the area surveyed, (**C**) average mitochondrial area, (**D**) the total mitochondrial area content per fiber area, and (**E**) the circularity index, which measures mitochondrial shape. For cristae, (**F**) the number of cristae (**G**) and cristae score, a measurement of the quality of cristae observed, are shown. Representative transmission electron micrographs for cardiac muscle at (**H**) 3-months, (**H’**) 1-year (**H’’**) and 2-years in female mice. Quantification of (**I**) the number of mitochondria normalized to the area surveyed, (**J**) average mitochondrial area, (**K**) mitochondrial area per fiber area, and (**L**) circularity index, (**M**) the number of cristae (**N**) and cristae score. (**O**) Schematic showing dysfunction of mitochondrial cristae across the aging process. Each dot represents a single mitochondrion. One-way ANOVA statistical test performed with post hoc Tukey’s Test. Significance values *, **, ***, **** indicate *p* ≤ 0.05, *p* ≤ 0.01, *p* ≤ 0.001, and *p* ≤ 0.0001, respectively.

Previous research has indicated that male mice have impaired mitochondrial function in cardiac muscle in comparison to female mice, in both healthy and cardiac pathological mice (41). To see mitochondrial and cristae structure changes across aging in a sex-dependent manner, we first looked at mitochondrial quantity which increased after a year of aging, while mitochondrial area was inversely proportional (Figure 1I-K). Like the male model, the circularity index of mitochondria increased with age (Figure 1L). Importantly, the female model recapitulated findings of significant loss of cristae across the aging process in murine cardiac tissue. Specifically, the number of cristae decreased (Figure 1M) consistently across aging, while the cristae score also significantly decreased past the 3-month age point (Figure 1N). Although in the female model, the cristae score improved slightly in comparing 2-year mice to 1-year mice, in both cohorts the average cristae score remained below 2, representing that 25% or more of mitochondrial area lacked typical cristae. Based on these findings, in tandem, we propose that mitochondrial fission, resulting in decreased mitochondrial area, is increased while the quality of cristae is decreased with age in a non-sex-dependent manner (Figure 1O). However, it should be noted that TEM can be limited in analyzing mitochondrial changes beyond those of cristae structure changes across the aging process. Given these changes did not present as sex-dependent, to properly understand how potential novel phenotypes arose in cardiac tissue, alterations in mitochondrial morphology were analyzed with 3D reconstruction in a male mouse model.

### Aging Changes Mitochondrial Size in Cardiac Muscle: 3D Reconstruction Analysis

Based on our observations of the lack of cristae folding in aging cardiac muscle, we employed 3D techniques to image cardiac muscle biopsies from young (3-month-old), mature (1-year-old), and aged (2-year-old) mice with SBF-SEM. With resolutions of 10 µm for the x- and y-planes and 50 µm for the z-plane, SBF-SEM enables 3D reconstruction of mitochondria with an accurate spatial resolution that cannot be seen in 2D. Given TEM did not reveal a sex-specific change, we focused on a male model. Specifically, we examined the morphological changes in the intermyofibrillar region as mitochondrial frequency has been demonstrated to increase in this region (42). Thus, there is a need to determine if such changes in frequency are associated with other mitochondrial-related aging changes, including mitochondrial orientation, network, and nanotunnel alterations. To elucidate the changes in intermyofibrillar mitochondria in relation to aging, we surveyed approximately 250 intermyofibrillar mitochondria from three male mice (n=3) (Figure 2A) sampled at each age time point with SBF-SEM 3D reconstruction methods. At a 10 µm by 10 µm image stack resolution, ∼50 50-µm ortho slides (Figure 2B) were manually traced at the transverse intervals (Figure 2C). This allowed for 3D reconstructions of mitochondria to be created, as observed in the flowchart of Figure 2 (Figure 2D). In totality, across the 3 mice, approximately 750 mitochondria for each age cohort were surveyed. All measurements used 3D metrics to understand the volumetric dynamics of mitochondria (Figure 2H).

**Figure 2:**
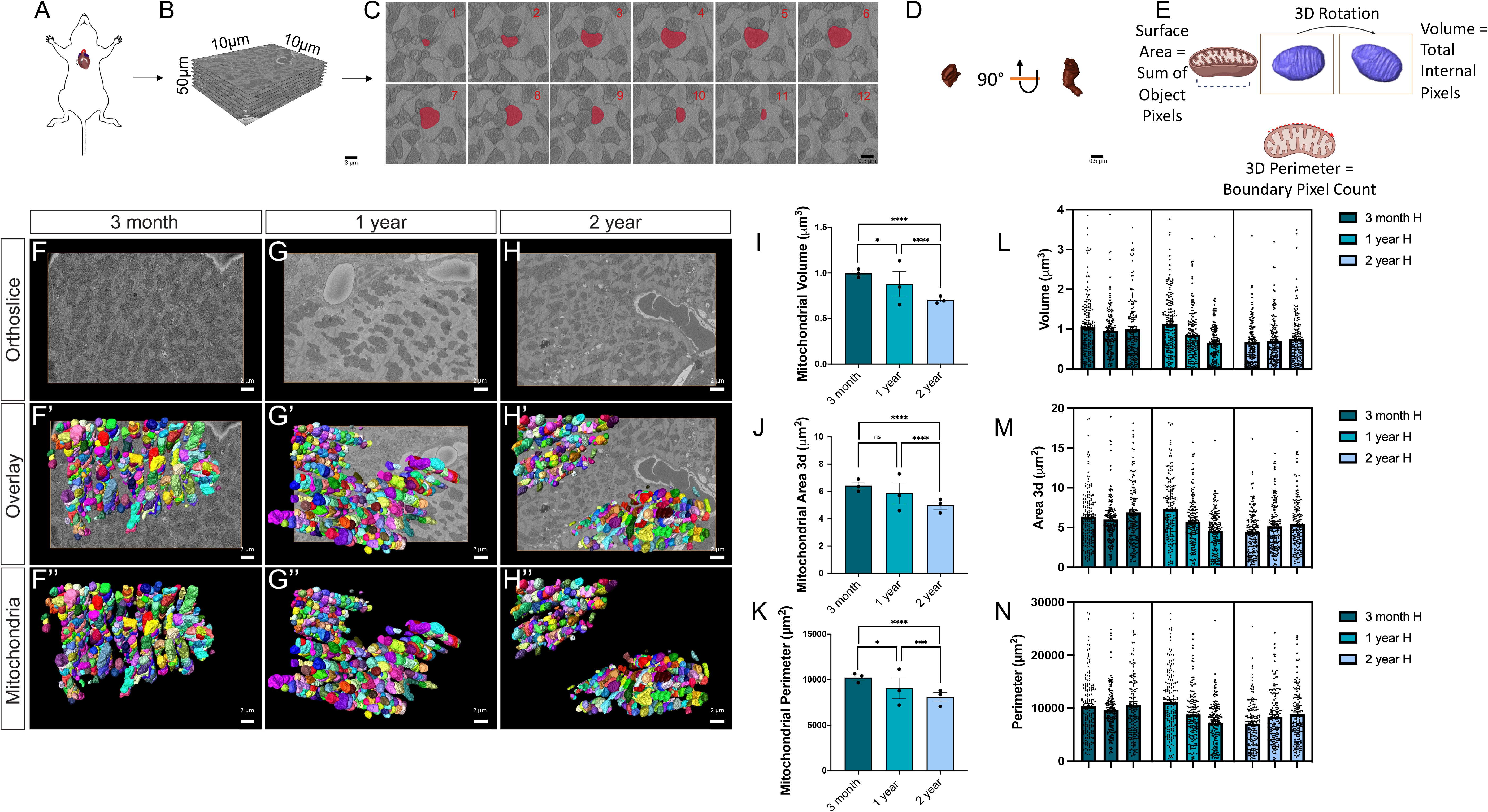
Changes in cardiac muscle mitochondria morphology across aging revealed in SBF-SEM. (**A**) Workflow depicting removal of the left ventricle of murine hearts. Following embedded fixation, SBF-SEM allows for (**B**) ortho slice alignment. (**C**) Manual segmentation of ortho slices was performed to (**D**) ultimately yield 3D reconstructions of mitochondria. (**E**) Schematic depicting how metrics are found using pre-installed analyses on Amira 3D Software. For some graphs, outlying dots were removed for presentation, but all mitochondria values were considered in statistical analysis. (**F-G**) Representative SBF-SEM orthoslice for cardiac muscle (n=3 mice). (**F’-G’**) 3D reconstructions of mitochondria in male cardiac tissues of different ages overlaid on ortho slices. (**F’’-G’’**) 3D reconstructed and isolated mitochondria for clear visualization. (**I-N**) 3D reconstructions were then quantified by (**I**) mitochondrial volume in cardiac muscle of different ages, (**J**) 3D area of the average mitochondria in cardiac muscle, and (**K**) Perimeter of the average mitochondria in cardiac muscle. Each dot represents the average of a single mouse. Individual quantifications for (**L**) average volume per mitochondria, (**M**) average 3D area of mitochondria, and (**N**) average perimeter of the mitochondria for each of the three mice sampled at 3-months, 1-year, and 2-years. One-way ANOVA statistical test performed with post hoc Tukey’s Test. Significance values *, **, ***, **** indicate *p* ≤ 0.05, *p* ≤ 0.01, *p* ≤ 0.001, and *p* ≤ 0.0001, respectively.

In Figure 2, we show representative images of the cardiac muscle at each aging point (Figure 2F-H). The overlay of the 3D reconstruction (Figure 2F’-H’), and the isolated 3D reconstruction (Figure 2F’’-H’’), allow for the mitochondrial structure to be viewed better (Video 1-2). Each color represents an independent mitochondrion. The perimeter, 3D area (or surface area), and volume decreased between 1-year to 2-year, which was a greater change than between 3-month and 1-year (Figure 2I-K). Importantly, this shows that while TEM was able to capture some dynamics of mitochondria, it neglected the alteration that is far greater in mice moving from adulthood to seniority (1-year to 2-year) than from adolescence to adulthood (3-month to 1-year). For these quantifications, three individual mice were sampled at each age (Figure 2L-N). When comparing the mitochondrial quantifications from each mouse, they showed overall little inter-group heterogeneity but persistent intra-individual variability.

With these observations, we sought to determine if mitochondrial networks changed in response to aging. To further elucidate age-related changes, we looked at cardiac muscle from both transverse (Figure 3A-C) and longitudinal views (Figure 3A’-C’). The mitochondrial branching index (MBI) was utilized to better understand mitochondrial branching (Figure 3D). The MBI measures network complexity by examining the ratio between transverse and longitudinal mitochondria (36) and is similar to aspect ratio which measures the ratio between major and minor axis (43). With this technique, we demonstrated that mitochondrial branching decreased between 3-month and the 1-year (Figure 3E). Similarly, within an age cohort, there is no significant variation between mice, but there was a large amount of heterogeneity in mitochondrial samples in each mouse (Figure 3F). To further characterize the mitochondrial types in each age cohort, we used mito-otyping, a method similar to karyotyping, to organize mitochondria based on their volumes to better visualize the overall mitochondrial diversity (Figure 3G). This allows for comparison of the mitochondria across ages at each volume. We measured sphericity to understand how the surface area changed during the aging process (Figure 3H). Sphericity generally showed minimal changes (Figure 3I) while there was homogeneity across all 9 mice measured (Figure 3J). For all of these metrics, some outliers were observed, which were omitted in the presentation for ease of view, but they were included in all statistical analysis (Figure S3). Critically, together, this approach revealed that there were few significant changes in morphology with only branching showing reductions. In combination, the aged cardiac muscle mitochondrial morphology resembled healthy mitochondria with a reduced size that lacks a phenotype or fragmentation (Figure 3). Since that the largest findings are loss in cristae structure and mitochondria size across aging, we sought to understand the underlying cause of these observed changes. Given the principal role of the MICOS complex in cristae formation and maintenance (27, 44), we investigated its role in cristae and mitochondrial remodeling across the aging process in cardiac muscle.

**Figure 3:**
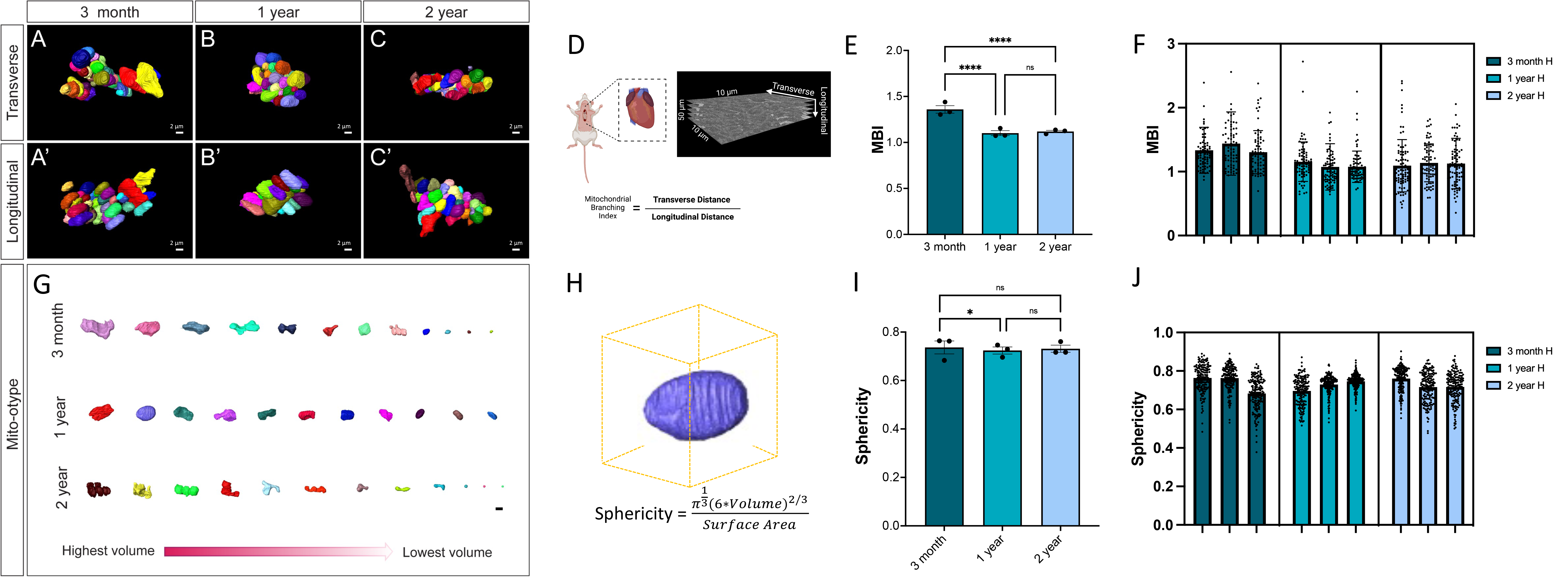
Changes in cardiac muscle branching and networking across aging revealed in SBF-SEM. (**A-C**) 3D reconstruction of individually colored mitochondria from a transverse view for mouse cardiac muscle of different ages. (**A’-C’**) 3D reconstruction of individually colored mitochondria from a longitudinal view in cardiac muscle tissues of different ages. (**D**) Schematic showing how transverse and longitudinal mitochondrial length is used to measure mitochondrial branching index. (**E**) Mitochondrial branching index was measured to estimate mitochondrial networks. (**F**) For each of the three male mice sampled at each aging time point, the mitochondrial branching index is shown. (**G**) Mitochondria 3D reconstructions were further organized by volume for each of the age cohorts. (**H**) Schematic showing how sphericity is measured, as a function of volume to surface area ratio. (**I**) To measure shape, sphericity changes across aging in cardiac muscle was further measured. (**J**) The average sphericity values from each of the approximately 75 mitochondria surveyed per mice is also shown. Outlying dots were removed for presentation for some graphs, but all mitochondria values were considered in statistical analysis. One-way ANOVA statistical test performed with post hoc Tukey’s Test. Significance value **** and ** indicates *p* ≤ 0.0001 and *p* ≤ 0.01.

### Aging Changes in MICOS Complex in Fibroblasts: 3D Reconstruction Analysis

Although it is established that the MICOS complex is critical for mitochondrial dynamics (22, 26), it is unclear how aging affects the MICOS complex. Studies have shown that *Opa1*, which is epistatic to the MICOS complex and physically interacts with components of the MICOS complex (23), decreases with age (45). With *Opa1* as a positive control, we sought to determine if the MICOS complex mRNA expression also decreased in cardiac muscle with age. As previously suggested (45), *Opa1* mRNA decreased by over 50% between 3-months and 2-years (Figure 4A). *Mitofilin* also decreased by 50% (Figure 4B). Likewise, *Chchd3* and *Chchd6* also progressively decreased with age but not as much as the decline of other transcripts (Figure 4C-D). While *Opa1* interacts with the MICOS complex, it is not required for the formation of cristae junctions at which the MICOS complex forms, nor does *Opa1* loss negatively affect MICOS components (46). This suggests that the loss of the MICOS complex across aging occurs in an *Opa1*-independent manner.

**Figure 4:**
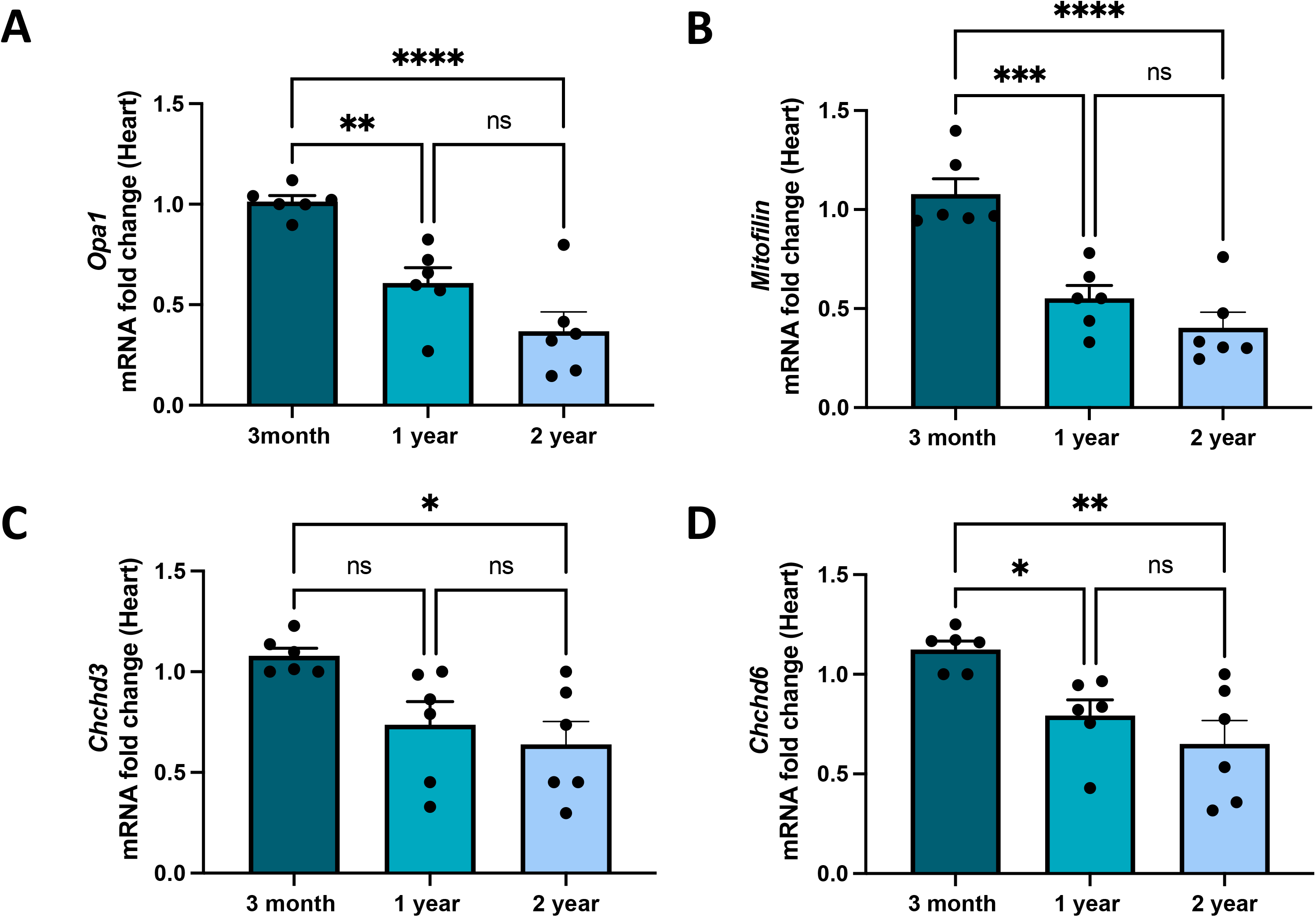
Transcription of *Opa-1* and MICOS genes in aging cardiac muscle. **(A-D)** qPCR analyzing the gene transcript fold changes of Opa-1 and MICOS across aging. (**A**) *Opa1* transcripts. (**B**) *Mitofilin* transcripts. (**C**) *Chchd3* transcripts. (**D**) *Chchd6* transcripts. One-way ANOVA statistical test performed with post hoc Tukey’s Test. Significance values *, **, ***, **** indicate *p* ≤ 0.05, *p* ≤ 0.01, *p* ≤ 0.001, and *p* ≤ 0.0001, respectively. For all experiments, n=6.

To further understand the role of mitochondrial dynamics upon the loss of these MICOS genes, we utilized CRISPR/Cas9 to make a knockout (KO) model in fibroblasts, which was validated by qPCR (Figure 5A-D). We have previously validated DRP1, MFN2, and OPA1 CRISPR-mediated gene knockout translates to protein level changes as shown by transmission electron microscopy (39, 47). Coupled with the current use of qPCR, gene knockout translates to reduced transcript and protein expression. Specifically, we measured approximately 1250 mitochondria across 10 cells. Since the loss of *Opa1* triggers changes in morphology (12, 15, 20, 39), we used it as a positive control for morphological changes. We marked mitochondria with MitoTracker Red and verified the successful deletion of *Opa1, Mitofilin, Chchd3,* and *Chchd6* (Figure 5E). Then z-stacks were reconstructed using Bitplane Imaris to quantify the number of cardiac fibroblasts with loss of MICOS expression (Figure 5E). Deletions of *Chchd3, Mitofilin, Chchd6,* or *Opa1* led to significant decreases in mitochondria length (Figure 5F) and volume (Figure 5G). We further sought to determine if the mitochondria were more tubular, which represents normal states, or fragmented, which represents stressed states (Figure 5H). Notably, *Chchd3-*KO cells had nearly completely fragmented mitochondria, like those in *Opa1-*KO. While the WT had mostly tubular mitochondria, the deletion of any of these critical mitochondrial proteins led to a much higher proportion of fragmented mitochondria. We further used live-confocal imaging to observe the dynamics of mitochondrial changes in real-time. The WT mitochondria (Video 3) were normal and potentially undergoing fusion. However, the cells with deletion of *Opa1* (Video 4), *Chchd3* (Video 5), or *Mitofilin* (Video 6) showed fragmented mitochondria. These videos aid in showing the real-time mitochondrial dynamics that may be contributing to changes in the 3D structure observed.

**Figure 5:**
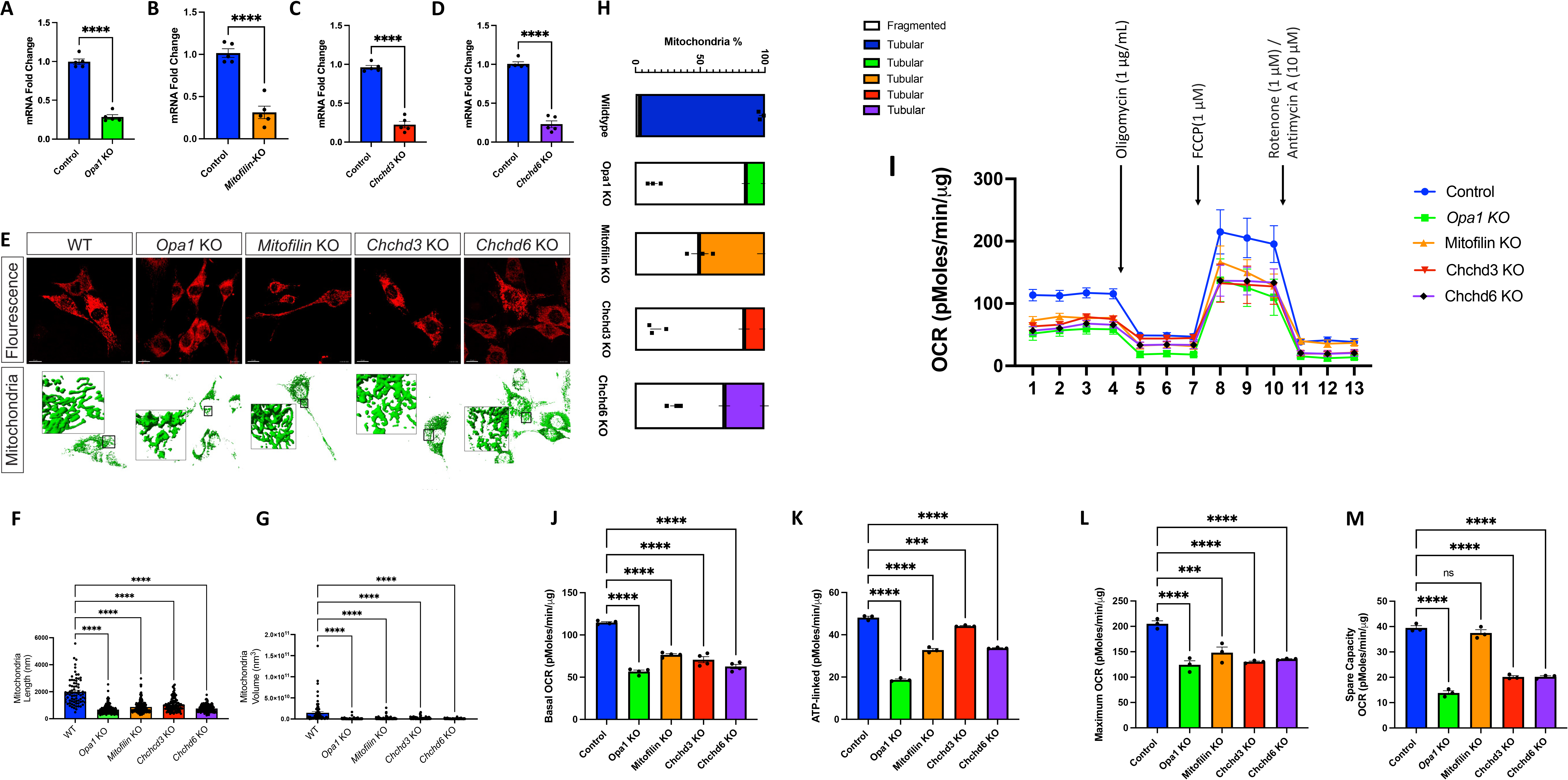
Loss of *Opa1* and MICOS genes result in mitochondrial structure changes and oxygen consumption rate changes. **(A)** qPCR (n=6) analyzing the gene transcript fold changes of Opa-1 and MICOS in cardiac fibroblasts upon CRISPR/Cas9 knockdown. (**A**) *Opa1* transcripts. (**B**) *Mitofilin* transcripts. (**C**) *Chchd3* transcripts. (**D**) *Chchd6* transcripts. **(E)** Confocal fluorescence (using mCherry-Mito-7) shows changes upon individual knockout (KO) of *Opa1, Mitofilin, Chchd3,* and *Chchd6* in mouse cardiac muscle. Below this, individual KO of *Opa1, Mitofilin, Chchd3,* and *Chchd6* was also observed in 3D reconstruction of SBF-SEM. (**F-H**) Quantification upon KO-state of each MICOS gene and *Opa1* (n=10 cells) was performed in 3D reconstruction. (**F**) Mitochondria length and (**G)** mitochondria volume across each knockout. (**H**) Additionally, the relative frequency of mitochondria that presented as fragmented or tubular was altered across knock outs. (**I**) Seahorse Analyzer was utilized to measure oxygen consumption rate (OCR) in *Opa1*, as positive control, along with *Chchd3*, *Chchd6,* and *Mitofilin* KD. (**J**) Basal OCR, which shows baseline energetic demands, (**K**) ATP-linked, which was measured by ATP-inhibitor oligomycin, (**L**) maximum OCR, which represents maximum respiration capacity upon application of FCCP. Finally, the application of rotenone/antimycin A causes only non-mitochondrial respiration to remain, which allows for measurement of (**M**) spare respiratory capacity, which signifies the extra amount of ATP which can quickly be generated. For seahorse n=6 plates for experimental knockouts, while for control n=16. For A-D: Unpaired (independent) student’s parametric statistical t-test performed. For all other statistical tests, One-way ANOVA statistical test performed with Dunnett’s multiple comparisons test.

To understand how the loss of the MICOS complex affects mitochondrial activity, we measured oxygen consumption rates (OCR) with an XF24 extracellular flux bioanalyzer upon knockout of *Opa1* and the MICOS genes in cardiac fibroblasts (Figure 5I). The OCR measurements encompassed four stages: basal respiration, ATP-linked respiration, maximal respiratory capacity, and reserve capacity. Basal respiration reflects the cellular OCR under normal, non-stressed conditions, while ATP-linked respiration represents the OCR directly related to ATP synthesis, which can be measured following the application of oligomycin, an inhibitor of ATP synthase. Maximal respiratory capacity refers to the maximum OCR a cell can achieve under stress or high-energy demand and is measured following the application of carbonyl cyanide-p-trifluoromethoxyphenylhydrazone (FCCP), an uncoupler of mitochondrial oxidative phosphorylation. Reserve capacity represents the difference between maximal respiratory capacity and basal respiration, indicating the cell’s ability to respond to increased energy demand and is measured following the application of Antimycin A/rotenone, inhibitors of the electron transport chain. The *Opa1*-KO, *Chchd3*-KO, *Chchd6*-KO, and *Mitofilin-*KO cells showed decreased basal OCR, ATP-linked OCR, and maximum OCR compared with the control (Figure 5J-M), indicating a general loss of OCR at baseline and during high demand. These results suggest that the depletion of the MICOS complex can lead to the impairment of the electron transfer chain, mitochondrial respiration, and bioenergetics, providing valuable insights for the development of targeted therapies for diseases associated with mitochondrial dysfunction. Of relevance, while *Opa1*-KO had a dramatic decrease in the reserve capacity of OCR, *Mitofilin*-KO cells showed no alteration in reserve capacity. These data show decreased oxygen consumption rate at basal consumption, decreased ATP-linked respiration following oligomycin application, decreased maximal respiratory capacity following FCCP application, and decreased reserve capacity, as measured following AntimycinA/rotenone treatment.

Loss of OPA1 does not yield changes in mtDNA (14), while alterations in the MICOS complex can affect mtDNA content (48, 49). Given these differential changes in mtDNA content, as shown previously, total protein is the most appropriate way to normalize to mitochondrial content (14, 50). Accordingly, cells were plated at a density of 20 × 10^3^ consistently with normalized protein, which together will take into account changes from the mitochondrial mass. Together this shows that changes in oxidative consumption occur independently from changes in mitochondrial mass. Overall, depletion of these critical mitochondrial proteins led to a general loss of oxygen consumption rate both at baseline and during high demand. These findings suggest that depletion of the MICOS complex can lead to the impairment of the electron transfer chain along with mitochondrial respiration and bioenergetics.

### Loss of *Chchd6* Results in Changes in Mitochondrial Size, Shape, and Fluorescence of Reactive Oxygen Species in Induced Pluripotent Stem Cell-Derived Cardiomyocytes

To understand the impact of the MICOS complex beyond murine cardiac fibroblast models, we knocked down *Chchd6*, a component of the MICOS complex, in induced pluripotent stem cell-derived cardiomyocytes (iPCS-CMs). To verify changes in mitochondrial size and morphology, we analyzed iPCS-CM through TEM (Figure 6A-B). We found that *Chchd6*-deficient iPCS-CM had reduced mitochondrial area and perimeter while mitochondrial circularity index increased (Figure 6C-F). We then moved to fluorescence live-cell imaging and overlaid tetramethylrhodamine, ethyl ester (TMRE) red dye, a label of active mitochondria and their membrane potential, dichlorodihydrofluorescein diacetate (DCFDA) green dye to assay for overall reactive oxygen species, and Hoechst blue dye to label for DNA in control iPCS-CMs (Figure 6G-G’’’) and *Chchd6*-depleted IPCS-CMs (Figure 6H-H’’’). Analysis showed that while mitochondrial membrane potential did not significantly change (Figure 6I), general reactive oxygen species increased (Figure 6J). Following verifying the cardiomyocyte identity of our iPSC-CM using troponin T staining (Figure 6K), to further validate this response and to look at alterations in levels of superoxides (51) upon loss of the MICOS complex, we examined Mitosox live-staining (Figure 6L-L’), which shows a significant increase of Mitosox fluorescence in the *Chchd6*-depleted, suggesting that the loss of this subunit of the MICOS complex results in increased in mitochondrial superoxides.

**Figure 6:**
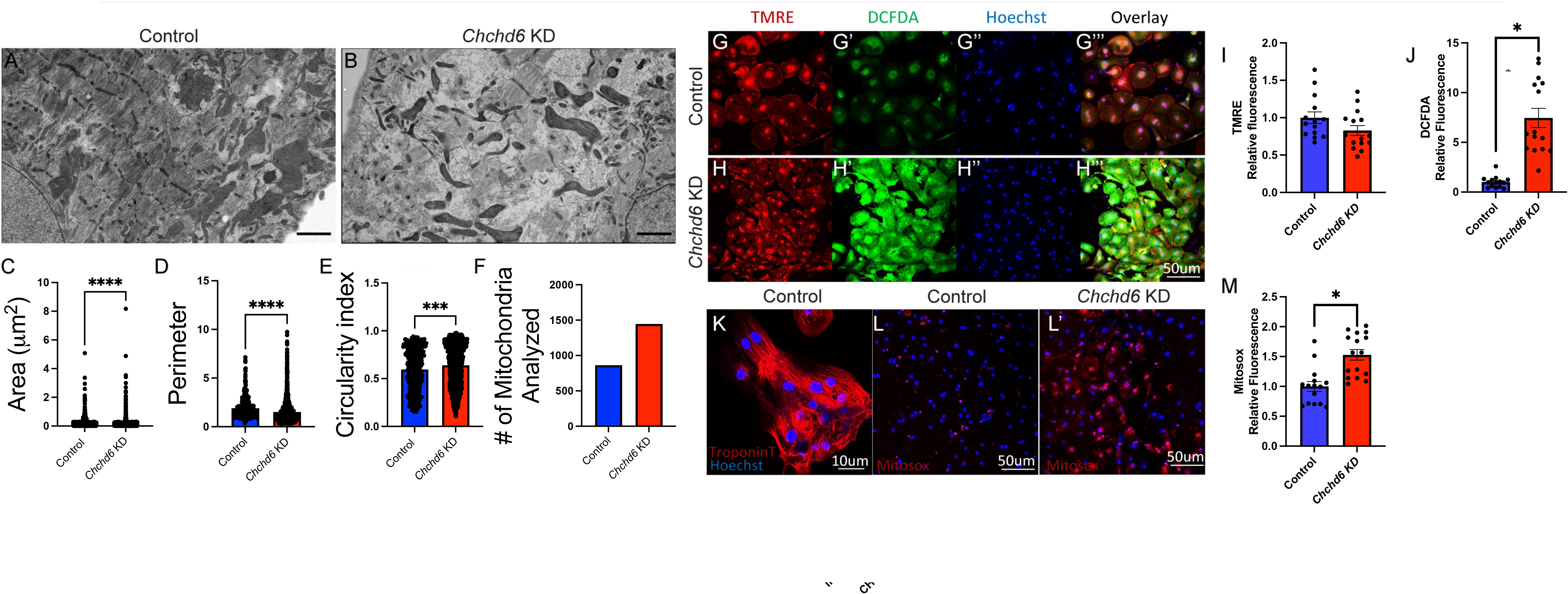
Chchd6-deficiency results in changes in mitochondrial amounts and reactive oxygen species in induced pluripotent stem cell-derived cardiomyocytes (iPSC-CMs). (**A**) Transmission electron microscopy (TEM) representative images of control and (**B**) CHCHD6-deficient induced pluripotent stem cell-derived cardiomyocytes. TEM analysis of (**C**) area, (**D**) perimeter, (**E**) circularity index, and (**F**) mitochondrial count (normalized to cell area). Fluorescence live-cell imaging in control iPSC-CMs used (**G**) tetramethylrhodamine, ethyl ester (TMRE; red) dye to label active mitochondria, (**G’**) Dichlorodihydrofluorescein diacetate (DCFDA; green) to assay for overall reactive oxygen species, (**G’’**) and Hoechst dye (blue) to label for DNA. Overlaid fluorescence image is also presented (**G’’’**) and fluorescence dye analysis were further performed in (**H-H’’’**) Chchd6-deficient IPSC-SMs. Analysis of fluorescent channels showing relative fluorescence of (**I**) TMRE and (**J**) DCFDA. Further fluorescence dye with Hoechst for DNA and Troponin-T (**K**) and MitoSox to measure reactive oxygen species including superoxides in control (**L**) and (**L’**) Chchd6-deficient IPSC-SMs. (**M**) Analysis of MitoSox fluorescence. (**C-F**) Each dot represents a single mitochondrion. (**I,J,M**) Each dot represents quantification from a microscopic field from three independent replicates for each control and KO groups. Unpaired (independent) student’s parametric statistical t-test performed. Significance value **** and * indicates *p* ≤ 0.0001 and *p* ≤ 0.05.

We also sought to understand the functional impact of loss of Chchd6 using the Vanderbilt BioVU bank (Figure S4A). This is a powerful biobank of over 270,000 collected by the Vanderbilt University Medical Center (52). This database includes a mixture of individuals of European ancestry and African American ancestry (Figure S4B). Genetically regulated gene expression (GREX) of CHCHD6 was calculated across cardiac tissues in individuals of European ancestry. The relationship between CHCHD6 GREX and the “Heart failure with reduced EF [Systolic or combined heart failure]” phenotype was evaluated across cases and controls within BioVU. The associations between CHCHD6 GREX and heart failure was nominally significant in 2 of the 3 tissues tested in individuals of European ancestry and 1 of the 3 tissues tested in individuals of African ancestry (p < 0.05). None of the associations met the across-tissue Bonferroni-corrected p-value (0.0167). Cases were required to have two instances of the “Heart failure with reduced EF [Systolic or combined heart failure]” phecode, controls had no reports of “Heart failure with reduced EF [Systolic or combined heart failure]” in their records. Individuals with similar codes were excluded from the analysis (Congestive heart failure; non-hypertensive, Congestive heart failure (CHF) NOS, Heart failure NOS, Heart failure with preserved EF [Diastolic heart failure], Ill-defined descriptions and complications of heart disease, Heart transplant/surgery, Abnormal function study of cardiovascular system, Symptoms involving cardiovascular system, Cardiac complications, not elsewhere classified). Covariates included sex, age, median age of medical record, genotype batch, and genetic ancestry (principle components 1-10). Together, this suggests that the MICOS complex may be linked to genetic factors of heart failure, and plays a role in modulating membrane potential and oxidative stress in iPCS-CM.

## DISCUSSION

Heart failure may be intrinsically linked to mitochondria (2, 33, 53), and understanding the interplay of cristae dynamics, MICOS complex component expression, and mitochondrial dynamics during aging may aid in the development of future therapies to restore energetic production. Here, we demonstrated abnormal mitochondrial structure and impaired function related to the loss of the MICOS complex. To our knowledge, this study is the first to show that abnormal MICOS complex alters mitochondrial morphology in cardiac tissue assessed by 3D-EM. Previous studies looking at mitochondria in aged cardiomyocytes did not find significant changes in mitochondrial number, however, there was no comprehensive quantitative analysis of the mitochondrial size and morphology (54). With the innovative 3D-EM technology, we reported decreased mitochondrial volume and altered morphology with age. Importantly, our study combined TEM and SBF-SEM to create novel 3D reconstructions of aging cardiac muscle cells. Although the high resolution of TEM allows us to measure cristae (39), it only creates 2D images that do not provide accurate spatial resolutions. Therefore, the 3D reconstructions of mitochondria by SBF-SEM allow us to better understand mitochondrial morphology. Future studies are needed that use focused ion beam-scanning electron microscopy (FIB-SEM) and thus, allow for better resolution of the details of smaller subcellular objects, such as cristae (36, 55). In the past, FIB-SEM was successfully performed in mouse cardiac muscle to characterize mitochondria, but not sub-organellar structures, such as cristae (54, 56). However, mixed-microscope approaches may still be required to achieve the resolution and sample size to analyze changes in mitochondria associated with aging and organ, particularly muscle, behavior.

Previous studies have quantified mitochondria with SBF-SEM in human and mouse muscles, yet the aging heart remains understudied with 3D techniques(36). Given that cardiac muscle is responsible for proper heart contractions, there remains a gap in our understanding of the changes of cardiac mitochondrial 3D structure in aging. When only examining TEM, our results show that much of the age-dependent changes in mitochondrial architecture occurs only from 3-months to 1-year, which is equivalent from adolescence to adult. However, when switching to 3D reconstruction, we observe that mitochondrial size and morphology progressively decrease across the aging process, including from 1-year to 2-years, which better emulates geriatric aging. Interestingly, however, mitochondrial networking does not appear as affected by the aging process past 1 year. In tandem, our 3D reconstruction elucidates how cardiac muscle mitochondria change across aging in mice, highlighting the importance of the MICOS complex in aging and its effects on cristae morphology and mitochondrial dynamics. Future studies are needed that examine mtDNA and other modulators of cristae and mitochondrial dynamics (57) as they may provide a further understanding of the MICOS complex in relation to mitochondrial structure.

When viewing mitochondrial 3D structure across aging, we looked for the presence of nanotunnels. Previous studies suggest that the formation of nanotunnels occurs during mitochondrial stress. Nanotunnels are mitochondrial structures that allow for the transport of materials and inter-mitochondrial communication (35). Nanotunnels have also been found in human skeletal muscles and in diseased states (35). We did not find evidence of nanotunnel formation in aged heart. This warrants future studies to investigate the factors leading to nanotunnel formation and whether this mechanism also exists in cardiac muscle (58).

We observed a decline in MICOS components gene expression with age (Figure 7). Additionally, with 3D reconstructions, we observed that the loss of *Opa1, Mitofilin, Chchd3,* and *Chchd6* in cardiac fibroblasts may be correlated with high mitochondrial dysfunction. This was marked by smaller mitochondria, altered morphology, and reduced OCR. Given the higher percentage of fragmented mitochondria and reduced mitochondrial volume of *Opa1*-KO, *Chchd3*-KO, *Chchd6*-KO, and *Mitofilin*-KO. The exact cause of this fragmentation is unclear, although past studies have linked cristae dysfunction and mitochondrial fragmentation, potentially through mtDNA mutations (25). However, further study is needed to determine if the loss of the MICOS complex is causing mitochondrial stress-related fragmentation, modulation of fission-fusion dynamics, or acts on mitochondrial dynamics in an altogether different way.

**Figure 7:**
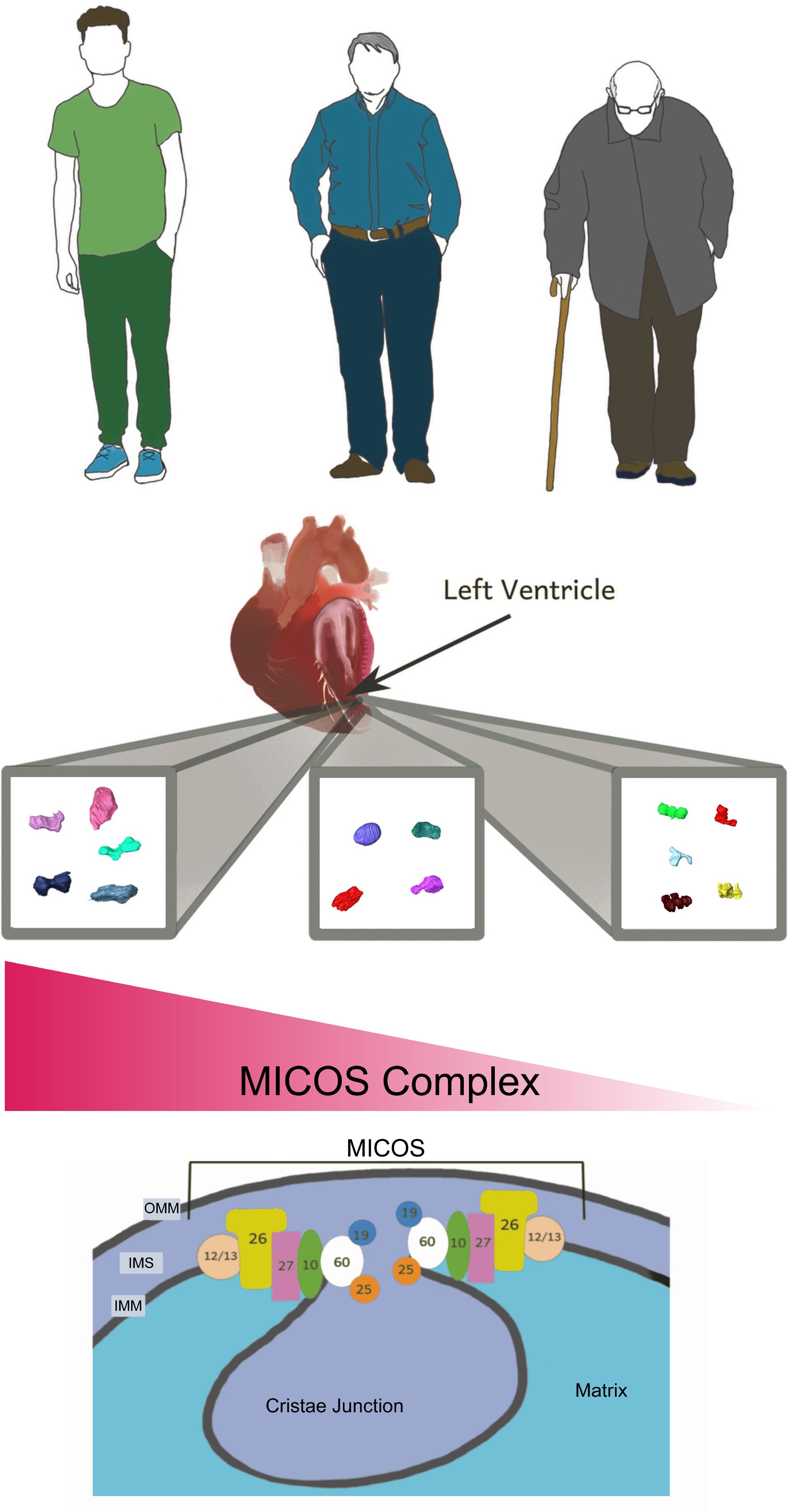
Graphical representation of loss of MICOS complex across aging that may occur, affecting the structure of mitochondria. Importantly, here we show this in a murine model, which may not translate to a human model (See Limitations section), but these findings suggest an important relation between the MICOS complex and aging which may be reproduced in human models.

It was predicted that these phenotypes would also arise in aging as expression of *Opa1*, *Mitofilin*, *Chchd3*, and *Chchd6* is reduced across aging (Figure 4). In cardiac tissue, we observed that the mitochondrial number increased with age, while the area decreased (Figure 1B-D). It is possible that fission rates may be increasing over time, due to pathway changes such as *Opa1* loss or *Drp1* upregulation (5, 20). Furthermore, the loss of cristae morphology supports the occurrence of age-associated-decline of the MICOS complex. When looking at 3D reconstruction, we principally noticed alterations in mitochondrial volume across aging which parallel findings from the loss of MICOS complex. In combination, these findings suggest that the loss of MICOS complex genes may be associated with a loss of overall mitochondrial volume. However, in comparing morphological changes in aged samples with MICOS complex KO fibroblasts, we noticed a differential change in morphology. Mitochondrial morphology in the heart did not change as drastically as anticipated in response to aging. In general, the sphericity of mitochondria did not undergo significant changes (Figure 3I).

Changes in mitochondrial shape are implicated in heart failure (33), which makes the specific study of cardiac muscle across aging essential for the development of new therapeutics. This study showed less networking, as measured through MBI, in cardiac mitochondria across aging than other studies have suggested (Figure 3). It is possible that some cardiac metabolic pathways are more resilient to aging changes, supporting the retention of mitochondrial morphology. Interestingly, a recent study showed that spermidine treatment restores mitochondrial morphology in the aged hearts (54), potentially indicating that spermidine differentially is regulated specifically in cardiac tissue. However, future studies are warranted to examine age-dependent mitochondrial changes in other tissues, to better understand tissue-specific mitochondrial changes that occur during aging.

Looking at iPCS-CMs, the loss of the MICOS complex led to increased ROS (Figure 6), consistent with previous studies in other models showing that loss of the MICOS complex may cause oxidative stress (59). Although the mitochondrial free radical theory of aging has been proposed for decades (60), the detailed mechanistic interplay between ROS and mitochondrial dynamics remains incompletely understood. Previous studies suggest that a positive feed loop within mitochondria promotes more ROS and thus, amplifies the ROS-related damage and structural decline (61). Consistent with this, loss of *Opa1* has been reported to cause the accumulation of ROS (62). Taken together, it is possible that during the aging process, the loss of *Opa1* and the MICOS complex allows for more oxidative stress to occur, resulting in ROS-induced ROS and the age-related functional decline of mitochondria (63). TMRE we used here to validate TEM findings that there are no significant changes in active mitochondria, thus oxidative stress changes can be attributed mainly due to altered generation (64). Interestingly, we also noticed that when using TMRE as a baseline measurement of mitochondrial membrane potential, we diverge from the literature in showing there is no change in the membrane potential. Loss of the MICOS complex, specifically Mic60, results in a decrease in membrane potential in Drosophila (48). A loss of membrane potential in OPA1 loss has also been seen in a murine model (14). Yet, TMRE may not be the best monitor of membrane potential as loss of negative charge affects its retention; thus, future studies may better investigate the MICOS-dependent loss of membrane potential (64, 66). Using the BioVU BioBank, we also observed linkages of *Chchd6* to factors of heart failure, with a differential ethnicity association (Figure S4), suggesting that loss of the MICOS complex and concomitant modulation of mitochondria structure has important implications in human health which must need to be further elucidated.

In summary, we combined TEM and 3D reconstructions to evaluate mitochondria and cristae morphology in male murine cardiac muscle. Our findings indicate that the MICOS complex decreases with age, and loss of the MICOS complex results in mitochondrial dysfunction and changes in mitochondrial structure. These findings better our understanding of how mitochondrial structure quantitively changes in cardiac muscle, and aid in understanding a potential that can be targeted to protect cardiac muscle mitochondria from complete fragmentation. Future studies are needed to explore this mechanism and continue to elucidate the link between age-related changes in cardiac cristae structural integrity, the MICOS complex, mitochondrial dysfunction, and ROS.

## LIMITATIONS

While we explored mitochondrial structure of around 750 mitochondria in each age cohort of male murine myocardium tissue, there are several potential limitations of our study. While the overall mitochondrial count was high, the quantity of mice in each age cohort is limited, although we did not note large heterogeneity between the mice for most mitochondrial quantifications. Notably, while Figure 7 shows a human model, this study was performed in a male murine model. There are several known critical differences between human and rodent hearts, which have differential developmental processes although they are generally considered to have similar anatomy (65). For example, while human hearts are predominantly composed of beta-Myosin heavy chain (MHC), in murine hearts, alpha-MHC is relatively more abundant while pathophysiology and aging causes a shift toward beta-MHC (67, 68). Notably, myocardial performance is partially dependent on the relative quantities of these MHCs, suggesting that they may modulate mitochondrial structure and function (69). While mitochondrial-related cardiovascular pathophysiology parallels each other between these models, greater research is still necessary in potential model-dependent differences. Beyond this, while we show through TEM that there are no sex-related differences on mitochondria across aging (Figure 1), future studies may still consider performing 3D reconstruction of female mice. While it is commonly understood that men are more vulnerable to age-related cardiovascular pathologies, female hearts have been shown to have increased resilience towards oxidative stress (70), which suggests that there may be a sex-dependent response to ROS. Finally, while the study sheds light on some of the molecular mechanisms underlying age-related decline in mitochondrial function, it does not address potential interventions or treatments to prevent or reverse this decline. Future research may look at if restoring the MICOS complex can restore these age-related changes. Since we only show a loss of MICOS transcripts over aging, it remains to be elucidated if increasing MICOS protein amount may restore mitochondrial structure.

## EXPERIMENTAL PROCEDURES

### Animal Care and Maintenance

These protocols are previously described (14), and animals used in this study were cared for using standard procedures approved by The University of Iowa Animal Care and Use Committee (IACUC) that follow the National Institute of Health Guide for the Care and Use of Laboratory Animals recommendations. WT male C57Bl/6J mice were exclusively used in experiments. They were housed at 22°C on a 12-hour light, 12-hour dark cycle. They had free access to water and standard chow. Mice were grown to various ages as described in the main text. 5% isoflurane/95% oxygen was used to anesthetize mice.

### RNA Extraction and RT-qPCR

RNA was extracted from tissue using TRIzol (Invitrogen) and RNeasy kit (Qiagen Inc). The subsequent concentration of isolated RNA samples was measured using a NanoDrop 1000 (NanoDrop products, Wilmington, DE, USA) spectrophotometer at an absorbance of 260 nm and 280 nm. Reverse transcription was conducted on isolated RNA (∼1 µg) using a High-Capacity cDNA Reverse Transcription Kit (Applied Biosciences, Carlsbad, CA). SYBR Green (Life Technologies, Carlsbad, CA) (1) was used for real-time quantitative PCR (qPCR). Three samples for each qPCR (∼50 ng) were placed in a 384-well plate that subsequently underwent qPCR in the ABI Prism 7900HT instrument (Applied Biosystems) (Table 1)(14). The following qPCR conditions were used: 1 cycle at 95 °C for 10 min; 40 cycles of 95 °C for 15 s; 59 °C for 15 s, 72 °C for 30 s, and 78 °C for 10 s; 1 cycle at 95 °C for 15 s; 1 cycle of 60 °C for 15 s; and one cycle of 95 °C for 15 s. The results were normalized to glyceraldehyde-3-phosphate dehydrogenase (*GAPDH*); data are shown as fold changes.

**Table 1:**
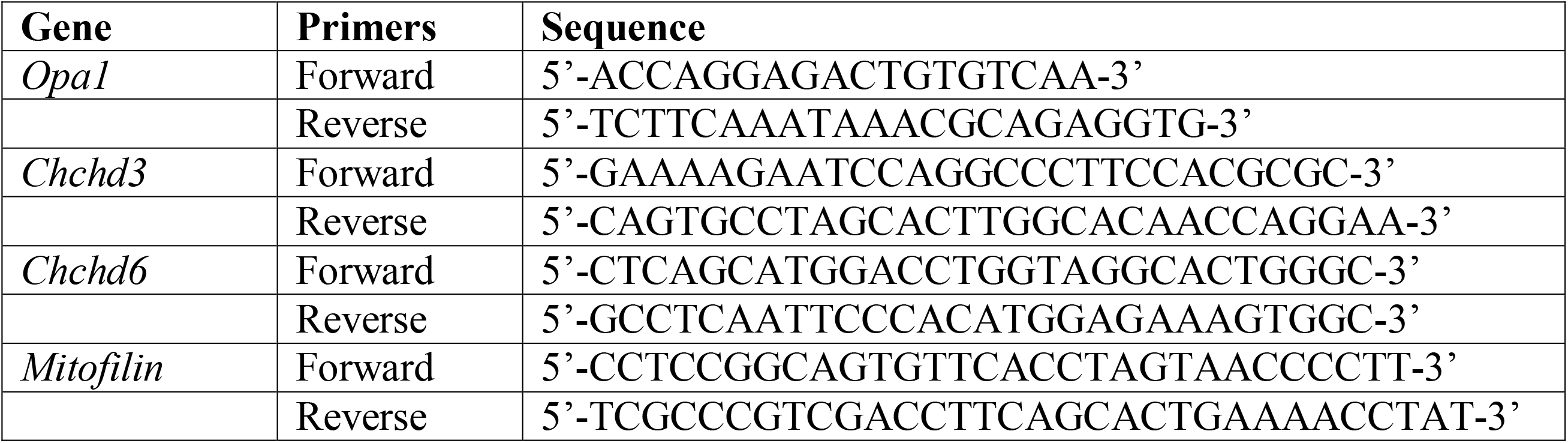
qPCR Primers Used.

### CRISPR-Cas9 Knockouts

Primary mouse fibroblasts were isolated as previously described (71, 72). Briefly, after cervical dislocation of 8-12 week mice, via sternotomy the heart was expose and transferred to 15 mL PBS centrifuge tube incubated at 37 °C. Aorta was removed and heart was washed with 10 mL of PBS warmed to 37 °C. Once ventricles were dissected, they minced into small pieces of 1 mm^3^, which were then subjected to enzymatic digestion using a mixture of Collagenase/Dispase and DNase I in PBS, while the tissues were incubated at 37 °C with 5% CO_2_ atmosphere for 10 min on a rocking platform. After being centrifuged for 5 minutes at 1500 rpm, the tissue digest pellet was resuspended in 1 mL of fibroblast culture medium (M199 medium supplemented with 10% FBS and 2% penicillin-streptomycin). This process was repeated approximately 9 times until the tissue was completely digested. Individual cell suspensions were combined pelleted again by running in a centrifuge at 1500 rpm for 7 minutes. Cells were resuspended in cardiac fibroblast media, and plated on 35 mm collagen-coated soft hydrogel-bound polystyrene plates. After the media is aspirated, plates with adherent cells were washed after 150 minutes with fibroblast culture medium. From there, 2 mL of fresh media was added, and the plate was incubated at 37 °C with 5% CO_2_ atmosphere. After 20 hours of digestion, this washing and replacing of media process was repeated, and cells were grown to 90% confluency.

Once isolated, CRISPR/Cas9 was used to infect cardiac fibroblasts to produce the following knockouts—control CRISPR/Cas9 (sc-418922), *Chchd6* (Mic25) CRISPR (sc-425817), *Chchd3* (Mic19) CRISPR (sc-425804), and *Mitofilin* (Mic60) CRISPR (sc-429376) (Santa Cruz Biotechnology, California, US) (Table 2). For each CRISPR, 2.5% of the CRISPR was combined with 2.5% RNAiMax (ThermoFisher Scientific; 13778075), and 95% Opti-MEM (Gibco; 31985070). This mixture was incubated at room temperature for 20 min. The media was removed from the cells and washing twice with PBS. 200 µL of the CRISPR mixture and 800 μL of Opti-MEM were added to each sample, then incubated at 37 °C for 4 hrs. An additional 1 mL of DMEM medium was added before cells were incubated at 37 °C overnight. Fibroblasts were washed with PBS, and a fresh medium was added. Experiments were performed 3- and 7-days following knockouts.

**Table 2:** Guide RNA and Plasmids Used.

| Gene Name | Type of Plasmid | CAS Number |
| --- | --- | --- |
| *Mitofilin* | CRISPR/Cas9 KO (m) | sc-429376 |
| *Chchd6* | CRISPR/Cas9 KO (m) | sc-425817 |
| *Chchd3* | CRISPR/Cas9 KO (m) | sc-425804 |
| *Control* | CRISPR/Cas9 KO (m) | sc-418922 |

### Processing of Mouse Left Ventricles for Serial Block-Face Scanning Electron Microscope (SBF-SEM)

SBF-SEM was performed as previously described (15). Male mice were euthanized using 5% isoflurane. The heart was excised and incubated in 2% glutaraldehyde with 100 mM phosphate buffer for 30 min. Left ventricles were dissected, cut into 1-mm^3^ cubes, and then incubated in 2.5% glutaraldehyde, 1% paraformaldehyde, and 120 mM sodium cacodylate solution for 1 hr.

From there, the tissue was three times washed with 100 mM cacodylate buffer at room temperature. The tissue was immersed in 3% potassium ferrocyanide and 2% osmium tetroxide for 1 hr at 4 °C, then washed three times with deionized water and incubated in 0.1% thiocarbohydrazide and 2% filtered osmium tetroxide 30 min. Finally, the tissue was washed three times with deionized water, before transferring to 1% uranyl acetate and left overnight at 4 °C. The next day, the samples were washed with deionized water, then incubated at 0.6% lead aspartate solution for 30 min at 60 °C. From there, the samples were dehydrated using an acetone gradient (20%, 50%, 70%, 90%, 95%, and 100% acetone for five mins each). Cardiac tissues were impregnated in Epoxy Taab 812 hard resin, then moved to new resin; polymerization occurred at 60°C for 36–48 hrs. Blocks of resin were sectioned, cut to 0.5 mm × 0.5 mm, and glued to aluminum pins. These pins were transferred to the FEI/Thermo Scientific Volumescope 2 SEM. 300−400 ultrathin (0.09 µm) serial sections from each block were collected for conventional TEM. All sections were collected onto formvar-coated slot grids (Pella, Redding CA), stained, and imaged.

### Measurement of OCR Using Seahorse

Oxygen consumption rate was measured for *Opa1*, *Cchchd3*, *Chchd6*, or *Mitofilin* KD fibroblasts using an XF24 bioanalyzer (Seahorse Bioscience: North Billerica, MA, USA), as previously described (14, 73). Fibroblasts were plated at a density of 20 × 10^3^ per well and differentiated for 3 days. For KO models, CRISPR/Cas9 for the relevant KO was added as per the protocol above prior to plating. The media was changed to XFLJDMEM (supplemented with 1 g/L D-Glucose, 0.11 g/L sodium pyruvate, and 4 mM L-Glutamine) and cells were incubated without CO_2_ for 60 minutes. oligomycin (1 μg/ml), carbonyl cyanide 4-(trifluoromethoxy)phenylhydrazone (FCCP; 1 μM), rotenone (1 μM), and antimycin A (10 μM (Figure 5I) was performed in that order while remaining in the XF-DMEM media. After measurement, cells were lysed accordingly to prior protocols (73), using 20 μl of 10 mM Tris with 0.1% Triton X-100 added at pH 7.4, and media replaced with 480 μl of Bradford reagent. Total protein concentration was then measured at absorbance at 595 nm and used for normalization. For each sample, three independent experiments were performed for each condition with representative data from the replicates being shown.

### Quantification of TEM Micrographs and Parameters Using ImageJ

The National Institutes of Health (NIH) *ImageJ* software was used to quantify TEM images, as previously described (39). Specifically, to ensure no bias in TEM quantification, one team member was responsible for conducting the experiment and fixing the cells and tissue, while another individual was responsible for processing and acquiring images using the electron microscope in a blinded and randomized manner. Each fiber of interest was divided into four equal quadrants, and two quadrants were randomly selected for measurements. Three additional blinded members were responsible for quantifying the anonymized samples. By averaging their collective findings, the potential for individual subjective bias was reduced. Moreover, those in charge of quantification were given randomized images at both whole-cell and higher magnification levels to further minimize bias.

### Segmentation and Quantification of 3D SBF-SEM Images Using Amira

SBF-SEM images were manually segmented in Amira to perform 3D reconstruction, as previously described (15). For each 3D reconstruction, (300−400 slices) were obtained and transferred to Amira. By hand, an individual blinded to samples, traced structural features manually on sequential slices of micrograph blocks. For each of the 3D reconstructions of cardiac muscle in mice, 50-100 serial sections were chosen at approximately equal z-direction intervals, stacked, aligned, and visualized using Amira to make videos and quantify volumetric structures. Algorithms for measurements were entered manually for those not already in the system (algorithms used are displayed in the main text; Figure 3D; Figure 3H). A total of 750 mitochondria from three mice were collected for each quantification.

### Confocal mCherry-Mito-7 Labeling

To label the mitochondria of cardiac fibroblasts, the mCherry-Mito-7 plasmid was transfected into the cells using a transfection reagent according to the manufacturer’s instructions (74). Briefly, the plasmid and the transfection reagent were diluted in Opti-MEM medium separately, mixed gently, and incubated for 20 min at room temperature. The mixture was added to the culture medium of the cells, and the cells were incubated for 24-48 h to allow expression of the mCherry-Mito-7 protein. The localization of the mCherry-Mito-7 protein in cardiac fibroblasts was visualized using a Leica SP8 Confocal Microscope. The cells were washed with PBS, fixed with 4% paraformaldehyde for 10 min, and mounted with DAPI-containing mounting medium. The fluorescent signals were observed using appropriate filters and recorded using a digital camera.

### Immunostaining protocol

For immunostaining, the hiPS-CMs were plated on glass-bottom dishes and were fixed in 4% paraformaldehyde for 15 minutes, permeabilized with 0.1% Triton-X-100 for 8 minutes and blocked with normal donkey serum for one hour, followed by incubation with primary antibody, Cardiac Troponin-T (Invitrogen, Catalog # MA5-12960) overnight. They were then probed with Donkey anti mouse AlexaFluor 555 secondary antibody (Thermo) and counterstained with DAPI. Images were acquired using a Leica SP8 confocal microscope followed by quantification using ImageJ software.

### Segmentation and Quantification of 3D Fibroblasts Using Confocal Microscopy and Imaris

For the 3D reconstruction of fibroblasts, a minimum of 10 cells were chosen, and approximately 20 mitochondria from each cell were segmented for a total of about 200 mitochondria. 3D structures were quantified as previously described (15) using the Imaris software (Bitplane), which automatically measured many parameters. All individuals measuring the parameters were blind to the experimental conditions but familiar in identifying diverse mitochondrial morphology. For each of the 3D reconstructions of cardiac fibroblasts, 50-100 serial sections were chosen at approximately equal z-direction intervals, stacked, aligned, and visualized.

### Maintenance of induced pluripotent stem cell-derived cardiomyocytes

Cultured on MatrigelLJcoated plates (Corning, Life Sciences), iPSC was fed 75% SFM and 25% mTeSR^+^. Cardiomyocyte differentiation was performed according to prior procedures (75), using STEMdiff™ Cardiomyocyte Differentiation and Maintenance kits. Typically, CMs began beating at day 8, and further enriched in day 10 of differentiation with lactic acid (sodium lLJlactate, Sigma–Aldrich, USA). To aid CM maturation, triiodothyronine (T3) was added across Days 20-40.

### Live-Cell Staining

Live cell staining procedures for iPSLJCMs were performed according to prior protocols (75). Once plated on glassLJbottom dishes, iPS-CMs media were replaced with 30 minutes with culture medium (37°C) supplemented with DCFDA (5 mM) and TMRE (25 nM). Media was replaced with new media supplemented with Hoechst 33342 for counterstaining. A Leica SP8 confocal microscope was used for image acquisition and ImageJ software was used for the analysis of fluorescence intensity.

### MitoSox

A stock solution of MitoSOXTM Red CMXRos-M7512 at a concentration of 1mM was prepared by adding 1 vial of MitoSOXTM Red (50µg) to 127µl of DMSO, which was diluted to a working solution of MitoSOXTM Red CMXRos at a concentration of 2.5µM by adding 10 µl of the stock solution to 10 ml of media. Cells were first aspirated of their media and then incubated with the working solution of MitoSOXTM Red at 37°C for 35 minutes. Following rinsing with warm media, cells were fixed with 16% paraformaldehyde at 37°C for 15 minutes. Following fixation, the cells were rinsed and imaged at 579 nm excitation and 599 nm emission wavelengths. Quantification of relative intensity normalized to standardized image area.

### Echocardiographic Parameters Collection

Animals were studied in accordance with protocols approved by the Institutional Animal Care and Use Committees of the Carver College of Medicine of the University of Iowa. Data from 9-10 animals at each age cohort were averaged to study in-vivo cardiac structure and function per previous procedures (76, 77). After anesthetization in an induction chamber using 2.5% isoflurane, mice were placed on a heated electrocardiography platform for heart rate monitoring during the imaging procedure and maintained at 37°C. Mice had a nose cone administering 1% isoflurane while in the left lateral decubitus position during imaging. A 13-MHz probe (Vivid V echocardiograph; GE Healthcare, Tampa, FL) was used to take standard B-mode M-mode images, along the short axis position of the papillary muscles.

### Data Analysis

All SBF-SEM and TEM data were obtained from at least three independent experiments and are presented as the mean across these experiments. In the presentation, black bars represent the standard error of mean, and dots represent individual data points shown. For all data with only two groups, an unpaired t-test was used. If more than two groups were compared, one-way ANOVA was performed, and significance was assessed using Tukey *post hoc* tests for multiple comparisons. For both analyses, GraphPad Prism software package was used (La Jolla, CA, USA). A minimum threshold of *p* < 0.05 indicated a significant difference. Higher degrees of statistical significance (**, ***, ****) were defined as *p* < 0.01, *p* < 0.001, and *p* < 0.0001, respectively. For genetically-regulated gene expression statistical analysis was performed as previously described (78). Briefly, transcriptome-wide association studies using gene, splicing, and proteome were compared with Bonferroni correction (0.05/#genes tested) for multiple testing correction. For all mouse models (Figure 2-3), each dot represents an independent mouse (n=3) but the statistical analysis shown represents the 750 mitochondria quantified.

## ACKNOWLEDGEMENTS

We would like to thank Mariya Sweetwyne for her advice as an aging expert.

---

**Figure S1.** Echocardiographic parameters of male and female mice at aged intervals of 3-month, 1-year, and 2-years. Presented as Means±SEM. (**A-I**) (**A**) HR indicates heart rate; (**B**) LVEF, left ventricle ejection fraction; (**C**) CO, cardiac output; (**D**) SV, stroke volume; (**E**) shows heart mass and (**F**) heart mass divided by body mass (**G**) LVT, left ventricle thickness; (**H**) ESV, end systolic volume; (**I**) EDV, end diastolic volume. Variable number of mice surveyed (n=9-10).

**Figure S2.** Changes in (**A**) body weight, (**B**) amount of lean mass, (**C**) amount of fat mass, and the mass of hearts (**D**) changes in murine mice across their aging. Each dot represents a mouse sampled. (**E**) Furthermore, glucose tolerance test shows there is altered glucose tolerance. (**F**) The area under curve of glucose tolerance test comparing the 3 age cohorts. Five mice surveyed (n=5). One-way ANOVA statistical test performed with post hoc Tukey’s Test.

**Figure S3.** Mitochondrial changes in (**A**) volume, (**B**) area 3D, (**C**) perimeter, (**D**) mitochondrial branching index, (**E**) and sphericity. All data are presented on a logarithmic scale.

**Figure S4.** (**A**) Genetically-regulated gene expression for CHCHD6 was calculated in BioVU participants using models built from the GTEx version 8 data (right panel), which contains genotype data matched to RNA-Seq data from 838 donors across 49 tissues. BioVU participants with a heart failure diagnosis were identified using ICD9/10 codes mapped to phecodes from Vanderbilt’s de-identified electronic health record database (middle panel). Imputed CHCHD6 expression was calculated and tested for association in heart failure cases versus controls using logistic regression models (right panel), accounting for genetic ancestry (principle components/PC 1-10), sex, age, and genotyping batch. (**B**) Demographics of BioVU Patients with Systolic Heart Failure, with standard deviations of any means in parenthesis (**C**) CHCHD6 GREX Association with Heart Failure Results in Cardiac Tissues. Beta level, standard error (SE), odds ratio of enrichment (OR), and p-value displayed for each tissue type.

**Video 1 –** 3D reconstruction of 3-month male murine cardiac tissue

**Video 2 --** 3D reconstruction of 2-year male murine cardiac tissue

**Video 3** – Confocal imaging of wildtype fibroblasts.

**Video 4** – Confocal imaging of *Opa1* KD fibroblasts.

**Video 5** – Confocal imaging of *Chchd3* KD fibroblasts.

**Video 6** – Confocal imaging of *Mitofilin* KD fibroblasts

## References

1. Boudina S, Sena S, Theobald H, Sheng X, Wright JJ, Hu XX, Aziz S, Johnson JI, Bugger H, Zaha VG, Abel ED. Mitochondrial energetics in the heart in obesity-related diabetes: direct evidence for increased uncoupled respiration and activation of uncoupling proteins. Diabetes 56: 2457–2466, 2007. doi: 10.2337/db07-0481.

2. Brown DA, Perry JB, Allen ME, Sabbah HN, Stauffer BL, Shaikh SR, Cleland JGF, Colucci WS, Butler J, Voors AA, Anker SD, Pitt B, Pieske B, Filippatos G, Greene SJ, Gheorghiade M. Mitochondrial function as a therapeutic target in heart failure. Nat Rev Cardiol 14: 238–250, 2017. doi: 10.1038/nrcardio.2016.203.

3. Tyumentsev MA, Stefanova NA, Kiseleva EV, Kolosova NG. Mitochondria with Morphology Characteristic for Alzheimer’s Disease Patients Are Found in the Brain of OXYS Rats. Biochemistry Moscow 83: 1083–1088, 2018. doi: 10.1134/S0006297918090109.

4. Alaimo A, Gorojod RM, Beauquis J, Muñoz MJ, Saravia F, Kotler ML. Deregulation of Mitochondria-Shaping Proteins Opa-1 and Drp-1 in Manganese-Induced Apoptosis. PLOS ONE 9: e91848, 2014. doi: 10.1371/journal.pone.0091848.

5. Lee Y, Jeong S-Y, Karbowski M, Smith CL, Youle RJ. Roles of the mammalian mitochondrial fission and fusion mediators Fis1, Drp1, and Opa1 in apoptosis. Molecular biology of the cell 15: 5001–5011, 2004.

6. Zuo W, Zhang S, Xia C-Y, Guo X-F, He W-B, Chen N-H. Mitochondria autophagy is induced after hypoxic/ischemic stress in a Drp1 dependent manner: the role of inhibition of Drp1 in ischemic brain damage. Neuropharmacology 86: 103–115, 2014.

7. Gunter TE, Yule DI, Gunter KK, Eliseev RA, Salter JD. Calcium and mitochondria. FEBS letters 567: 96–102, 2004.

8. Mehta MM, Weinberg SE, Chandel NS. Mitochondrial control of immunity: beyond ATP. Nat Rev Immunol 17: 608–620, 2017. doi: 10.1038/nri.2017.66.

9. Nicholls DG. Mitochondria and calcium signaling. Cell calcium 38: 311–317, 2005.

10. Barja G. The mitochondrial free radical theory of aging. Progress in molecular biology and translational science 127: 1–27, 2014.

11. Figueiredo PA, Mota MP, Appell HJ, Duarte JA. The role of mitochondria in aging of skeletal muscle. Biogerontology 9: 67–84, 2008. doi: 10.1007/s10522-007-9121-7.

12. Cipolat S, de Brito OM, Dal Zilio B, Scorrano L. OPA1 requires mitofusin 1 to promote mitochondrial fusion. Proceedings of the National Academy of Sciences 101: 15927–15932, 2004. doi: 10.1073/pnas.0407043101.

13. Otera H, Miyata N, Kuge O, Mihara K. Drp1-dependent mitochondrial fission via MiD49/51 is essential for apoptotic cristae remodeling. Journal of Cell Biology 212: 531– 544, 2016.

14. Pereira RO, Tadinada SM, Zasadny FM, Oliveira KJ, Pires KMP, Olvera A, Jeffers J, Souvenir R, Mcglauflin R, Seei A. OPA 1 deficiency promotes secretion of FGF 21 from muscle that prevents obesity and insulin resistance. The EMBO journal 36: 2126–2145, 2017.

15. Garza-Lopez E, Vue Z, Katti P, Neikirk K, Biete M, Lam J, Beasley HK, Marshall AG, Rodman TA, Christensen TA, Salisbury JL, Vang L, Mungai M, AshShareef S, Murray SA, Shao J, Streeter J, Glancy B, Pereira RO, Abel ED, Hinton A. Protocols for Generating Surfaces and Measuring 3D Organelle Morphology Using Amira. Cells 11: 65, 2022. doi: 10.3390/cells11010065.

16. Wang C, Du W, Su QP, Zhu M, Feng P, Li Y, Zhou Y, Mi N, Zhu Y, Jiang D, Zhang S, Zhang Z, Sun Y, Yu L. Dynamic tubulation of mitochondria drives mitochondrial network formation. Cell Res 25: 1108–1120, 2015. doi: 10.1038/cr.2015.89.

17. Glancy B, Kim Y, Katti P, Willingham TB. The Functional Impact of Mitochondrial Structure Across Subcellular Scales [Online]. Frontiers in Physiology 11, 2020. https://www.frontiersin.org/articles/10.3389/fphys.2020.541040 [16 Dec. 2022].

18. Ahmad T, Aggarwal K, Pattnaik B, Mukherjee S, Sethi T, Tiwari B, Kumar M, Micheal A, Mabalirajan U, Ghosh B, Roy S, Agrawal A. Computational classification of mitochondrial shapes reflects stress and redox state. Cell death & disease 4: e461, 2013. doi: 10.1038/cddis.2012.213.

19. Rambold AS, Kostelecky B, Elia N, Lippincott-Schwartz J. Tubular network formation protects mitochondria from autophagosomal degradation during nutrient starvation. Proc Natl Acad Sci U S A 108: 10190–10195, 2011. doi: 10.1073/pnas.1107402108.

20. Frezza C, Cipolat S, De Brito OM, Micaroni M, Beznoussenko GV, Rudka T, Bartoli D, Polishuck RS, Danial NN, De Strooper B. OPA1 controls apoptotic cristae remodeling independently from mitochondrial fusion. Cell 126: 177–189, 2006.

21. Cogliati S, Enriquez JA, Scorrano L. Mitochondrial cristae: where beauty meets functionality. Trends in biochemical sciences 41: 261–273, 2016.

22. Kozjak-Pavlovic V. The MICOS complex of human mitochondria. Cell and tissue research 367: 83–93, 2017.

23. Glytsou C, Calvo E, Cogliati S, Mehrotra A, Anastasia I, Rigoni G, Raimondi A, Shintani N, Loureiro M, Vazquez J. Optic atrophy 1 is epistatic to the core MICOS component MIC60 in mitochondrial cristae shape control. Cell reports 17: 3024–3034, 2016.

24. Eramo MJ, Lisnyak V, Formosa LE, Ryan MT. The ‘mitochondrial contact site and cristae organising system’ (MICOS) in health and human disease. The Journal of Biochemistry 167: 243–255, 2020. doi: 10.1093/jb/mvz111.

25. Kondadi AK, Anand R, Hänsch S, Urbach J, Zobel T, Wolf DM, Segawa M, Liesa M, Shirihai OS, WeidtkampOPeters S, Reichert AS. Cristae undergo continuous cycles of membrane remodelling in a MICOSLJdependent manner. EMBO Rep 21: e49776, 2020. doi: 10.15252/embr.201949776.

26. Friedman JR, Mourier A, Yamada J, McCaffery JM, Nunnari J. MICOS coordinates with respiratory complexes and lipids to establish mitochondrial inner membrane architecture. Elife 4: e07739, 2015.

27. Stephan T, Brüser C, Deckers M, Steyer AM, Balzarotti F, Barbot M, Behr TS, Heim G, Hübner W, Ilgen P. MICOS assembly controls mitochondrial inner membrane remodeling and crista junction redistribution to mediate cristae formation. The EMBO journal 39: e104105, 2020.

28. Zhang J, He Z, Fedorova J, Logan C, Bates L, Davitt K, Le V, Murphy J, Li M, Wang M, Lakatta EG, Ren D, Li J. Alterations in mitochondrial dynamics with age-related Sirtuin1/Sirtuin3 deficiency impair cardiomyocyte contractility. Aging Cell 20: e13419, 2021. doi: 10.1111/acel.13419.

29. Parra V, Verdejo HE, Iglewski M, del Campo A, Troncoso R, Jones D, Zhu Y, Kuzmicic J, Pennanen C, LopezOCrisosto C, Jaña F, Ferreira J, Noguera E, Chiong M, Bernlohr DA, Klip A, Hill JA, Rothermel BA, Abel ED, Zorzano A, Lavandero S. Insulin Stimulates Mitochondrial Fusion and Function in Cardiomyocytes via the Akt-mTOR-NFκB-Opa-1 Signaling Pathway. Diabetes 63: 75–88, 2013. doi: 10.2337/db13-0340.

30. Kim J, Wei Y, Sowers JR. Role of mitochondrial dysfunction in insulin resistance. Circulation research 102: 401–414, 2008.

31. Friederich M, Hansell P, Palm F. Diabetes, oxidative stress, nitric oxide and mitochondria function. Current diabetes reviews 5: 120–144, 2009.

32. Ngoh GA, Papanicolaou KN, Walsh K. Loss of mitofusin 2 promotes endoplasmic reticulum stress. J Biol Chem 287: 20321–20332, 2012. doi: 10.1074/jbc.M112.359174.

33. Kenny HC, Abel ED. Heart Failure in Type 2 Diabetes Mellitus. Circ Res 124: 121–141, 2019. doi: 10.1161/CIRCRESAHA.118.311371.

34. Willingham TB, Ajayi PT, Glancy B. Subcellular Specialization of Mitochondrial Form and Function in Skeletal Muscle Cells. Front Cell Dev Biol 9: 757305, 2021. doi: 10.3389/fcell.2021.757305.

35. Vincent AE, Turnbull DM, Eisner V, Hajnóczky G, Picard M. Mitochondrial Nanotunnels. Trends in Cell Biology 27: 787–799, 2017. doi: 10.1016/j.tcb.2017.08.009.

36. Vincent AE, White K, Davey T, Philips J, Ogden RT, Lawless C, Warren C, Hall MG, Ng YS, Falkous G, Holden T, Deehan D, Taylor RW, Turnbull DM, Picard M. Quantitative 3D Mapping of the Human Skeletal Muscle Mitochondrial Network. Cell Reports 26: 996-1009.e4, 2019. doi: 10.1016/j.celrep.2019.01.010.

37. Lesnefsky EJ, Chen Q, Hoppel CL. Mitochondrial Metabolism in Aging Heart. Circ Res 118: 1593–1611, 2016. doi: 10.1161/CIRCRESAHA.116.307505.

38. Chaudhary KR, El-Sikhry H, Seubert JM. Mitochondria and the aging heart. J Geriatr Cardiol 8: 159–167, 2011. doi: 10.3724/SP.J.1263.2011.00159.

39. Lam J, Katti P, Biete M, Mungai M, AshShareef S, Neikirk K, Garza Lopez E, Vue Z, Christensen TA, Beasley HK, Rodman TA, Murray SA, Salisbury JL, Glancy B, Shao J, Pereira RO, Abel ED, Hinton A. A Universal Approach to Analyzing Transmission Electron Microscopy with ImageJ. Cells 10: 2177, 2021. doi: 10.3390/cells10092177.

40. Eisner V, Picard M, Hajnóczky G. Mitochondrial dynamics in adaptive and maladaptive cellular stress responses. Nat Cell Biol 20: 755–765, 2018. doi: 10.1038/s41556-018-0133-0.

41. John C, Grune J, Ott C, Nowotny K, Deubel S, Kühne A, Schubert C, Kintscher U, Regitz-Zagrosek V, Grune T. Sex Differences in Cardiac Mitochondria in the New Zealand Obese Mouse [Online]. Frontiers in Endocrinology 9, 2018. https://www.frontiersin.org/articles/10.3389/fendo.2018.00732 [26 Jan. 2023].

42. Vendelin M, Béraud N, Guerrero K, Andrienko T, Kuznetsov AV, Olivares J, Kay L, Saks VA. Mitochondrial regular arrangement in muscle cells: a “crystal-like” pattern. American Journal of Physiology-Cell Physiology 288: C757–C767, 2005. doi: 10.1152/ajpcell.00281.2004.

43. Picard M, White K, Turnbull DM. Mitochondrial morphology, topology, and membrane interactions in skeletal muscle: a quantitative three-dimensional electron microscopy study. J Appl Physiol (1985) 114: 161–171, 2013. doi: 10.1152/japplphysiol.01096.2012.

44. Hu C, Shu L, Huang X, Yu J, Li L, Gong L, Yang M, Wu Z, Gao Z, Zhao Y, Chen L, Song Z. OPA1 and MICOS Regulate mitochondrial crista dynamics and formation. Cell Death Dis 11: 1–17, 2020. doi: 10.1038/s41419-020-03152-y.

45. Tezze C, Romanello V, Desbats MA, Fadini GP, Albiero M, Favaro G, Ciciliot S, Soriano ME, Morbidoni V, Cerqua C, Loefler S, Kern H, Franceschi C, Salvioli S, Conte M, Blaauw B, Zampieri S, Salviati L, Scorrano L, Sandri M. Age-Associated Loss of OPA1 in Muscle Impacts Muscle Mass, Metabolic Homeostasis, Systemic Inflammation, and Epithelial Senescence. Cell Metab 25: 1374-1389.e6, 2017. doi: 10.1016/j.cmet.2017.04.021.

46. Barrera M, Koob S, Dikov D, Vogel F, Reichert AS. OPA1 functionally interacts with MIC60 but is dispensable for crista junction formation. FEBS Letters 590: 3309–3322, 2016. doi: 10.1002/1873-3468.12384.

47. Hinton A, Katti P, Christensen TA, Mungai M, Shao J, Zhang L, Trushin S, Alghanem A, Jaspersen A, Geroux RE, Neikirk K, Biete M, Lopez EG, Shao B, Vue Z, Vang L, Beasley HK, Marshall AG, Stephens D, Damo S, Ponce J, Bleck CKE, Hicsasmaz I, Murray SA, Edmonds RAC, Dajles A, Koo YD, Bacevac S, Salisbury JL, Pereira RO, Glancy B, Trushina E, Abel ED. A Comprehensive Approach to Sample Preparation for Electron Microscopy and the Assessment of Mitochondrial Morphology in Tissue and Cultured Cells. .

48. Wang L, Hsu T, Lin H, Fu C. Drosophila MICOS knockdown impairs mitochondrial structure and function and promotes mitophagy in muscle tissue. Biology Open 9: bio054262, 2020. doi: 10.1242/bio.054262.

49. Qin J, Guo Y, Xue B, Shi P, Chen Y, Su QP, Hao H, Zhao S, Wu C, Yu L, Li D, Sun Y. ER-mitochondria contacts promote mtDNA nucleoids active transportation via mitochondrial dynamic tubulation. Nat Commun 11: 4471, 2020. doi: 10.1038/s41467-020-18202-4.

50. Pereira RO, Marti A, Olvera AC, Tadinada SM, Bjorkman SH, Weatherford ET, Morgan DA, Westphal M, Patel PH, Kirby AK, Hewezi R, Bùi Trân W, García-Peña LM, Souvenir RA, Mittal M, Adams CM, Rahmouni K, Potthoff MJ, Abel ED. OPA1 deletion in brown adipose tissue improves thermoregulation and systemic metabolism via FGF21. eLife 10: e66519, 2021. doi: 10.7554/eLife.66519.

51. Dikalov SI, Harrison DG. Methods for Detection of Mitochondrial and Cellular Reactive Oxygen Species. Antioxid Redox Signal 20: 372–382, 2014. doi: 10.1089/ars.2012.4886.

52. Davis L. Psychiatric Genomics, Phenomics, and Ethics Research In A 270,000-Person Biobank (BioVU). European Neuropsychopharmacology 29: S739–S740, 2019. doi: 10.1016/j.euroneuro.2017.06.069.

53. Bayeva M, Gheorghiade M, Ardehali H. Mitochondria as a Therapeutic Target in Heart Failure. Journal of the American College of Cardiology 61: 599–610, 2013. doi: 10.1016/j.jacc.2012.08.1021.

54. Messerer J, Wrede C, Schipke J, Brandenberger C, Abdellatif M, Eisenberg T, Madeo F, Sedej S, Mühlfeld C. Spermidine supplementation influences mitochondrial number and morphology in the heart of aged mice. Journal of Anatomy 242: 91–101, 2023. doi: 10.1111/joa.13618.

55. Baena V, Conrad R, Friday P, Fitzgerald E, Kim T, Bernbaum J, Berensmann H, Harned A, Nagashima K, Narayan K. FIB-SEM as a Volume Electron Microscopy Approach to Study Cellular Architectures in SARS-CoV-2 and Other Viral Infections: A Practical Primer for a Virologist. Viruses 13: 611, 2021. doi: 10.3390/v13040611.

56. Glancy B, Hartnell LM, Malide D, Yu Z-X, Combs CA, Connelly PS, Subramaniam S, Balaban RS. Mitochondrial reticulum for cellular energy distribution in muscle. Nature 523: 617–620, 2015.

57. Kondadi AK, Anand R, Reichert AS. Functional Interplay between Cristae Biogenesis, Mitochondrial Dynamics and Mitochondrial DNA Integrity. Int J Mol Sci 20: 4311, 2019. doi: 10.3390/ijms20174311.

58. Lavorato M, Iyer VR, Dewight W, Cupo RR, Debattisti V, Gomez L, Fuente SD la, Zhao Y-T, Valdivia HH, Hajnóczky G, Franzini-Armstrong C. Increased mitochondrial nanotunneling activity, induced by calcium imbalance, affects intermitochondrial matrix exchanges. PNAS 114: E849–E858, 2017. doi: 10.1073/pnas.1617788113.

59. Warnsmann V, Marschall L-M, Meeßen AC, Wolters M, Schürmanns L, Basoglu M, Eimer S, Osiewacz HD. Disruption of the MICOS complex leads to an aberrant cristae structure and an unexpected, pronounced lifespan extension in Podospora anserina. Journal of Cellular Biochemistry 123: 1306–1326, 2022. doi: 10.1002/jcb.30278.

60. Sanz A, Stefanatos RK. The mitochondrial free radical theory of aging: a critical view. Current aging science 1: 10–21, 2008.

61. Ježek J, Cooper KF, Strich R. Reactive Oxygen Species and Mitochondrial Dynamics: The Yin and Yang of Mitochondrial Dysfunction and Cancer Progression. Antioxidants 7, 2018. doi: 10.3390/antiox7010013.

62. Quintana-Cabrera R, Manjarrés-Raza I, Vicente-Gutiérrez C, Corrado M, Bolaños JP, Scorrano L. Opa1 relies on cristae preservation and ATP synthase to curtail reactive oxygen species accumulation in mitochondria. Redox Biol 41: 101944, 2021. doi: 10.1016/j.redox.2021.101944.

63. Haas RH. Mitochondrial Dysfunction in Aging and Diseases of Aging. Biology 8: 48, 2019. doi: 10.3390/biology8020048.

64. Chazotte B. Labeling mitochondria with TMRM or TMRE. Cold Spring Harb Protoc 2011: 895–897, 2011. doi: 10.1101/pdb.prot5641.

65. Wessels A, Sedmera D. Developmental anatomy of the heart: a tale of mice and man. Physiological Genomics 15: 165–176, 2003. doi: 10.1152/physiolgenomics.00033.2003.

66. Chazotte B. Labeling Mitochondria with MitoTracker Dyes. Cold Spring Harb Protoc 2011: pdb.prot5648, 2011. doi: 10.1101/pdb.prot5648.

67. Krenz M, Robbins J. Impact of beta-myosin heavy chain expression on cardiac function during stress. Journal of the American College of Cardiology 44: 2390–2397, 2004. doi: 10.1016/j.jacc.2004.09.044.

68. Carnes CA, Geisbuhler TP, Reiser PJ. Age-dependent changes in contraction and regional myocardial myosin heavy chain isoform expression in rats. Journal of Applied Physiology 97: 446–453, 2004. doi: 10.1152/japplphysiol.00439.2003.

69. Herron TJ, McDonald KS. Small Amounts of α-Myosin Heavy Chain Isoform Expression Significantly Increase Power Output of Rat Cardiac Myocyte Fragments. Circulation Research 90: 1150–1152, 2002. doi: 10.1161/01.RES.0000022879.57270.11.

70. Kander MC, Cui Y, Liu Z. Gender difference in oxidative stress: a new look at the mechanisms for cardiovascular diseases. Journal of Cellular and Molecular Medicine 21: 1024–1032, 2017. doi: 10.1111/jcmm.13038.

71. Gansemer ER, McCommis KS, Martino M, King-McAlpin AQ, Potthoff MJ, Finck BN, Taylor EB, Rutkowski DT. NADPH and Glutathione Redox Link TCA Cycle Activity to Endoplasmic Reticulum Homeostasis. iScience 23: 101116, 2020. doi: 10.1016/j.isci.2020.101116.

72. Sahadevan P, Allen BG. Isolation and culture of adult murine cardiac atrial and ventricular fibroblasts and myofibroblasts. Methods 203: 187–195, 2022. doi: 10.1016/j.ymeth.2021.04.004.

73. Dranka BP, Benavides GA, Diers AR, Giordano S, Zelickson BR, Reily C, Zou L, Chatham JC, Hill BG, Zhang J, Landar A, Darley-Usmar VM. Assessing bioenergetic function in response to oxidative stress by metabolic profiling. Free Radic Biol Med 51: 1621–1635, 2011. doi: 10.1016/j.freeradbiomed.2011.08.005.

74. Olenych SG, Claxton NS, Ottenberg GK, Davidson MW. The fluorescent protein color palette. Curr Protoc Cell Biol Chapter 21: Unit 21.5, 2007. doi: 10.1002/0471143030.cb2105s36.

75. Yoon J, Daneshgar N, Chu Y, Chen B, Hefti M, Vikram A, Irani K, Song L, Brenner C, Abel ED, London B, Dai D. Metabolic rescue ameliorates mitochondrial encephaloLJcardiomyopathy in murine and human iPSC models of Leigh syndrome. Clin Transl Med 12: e954, 2022. doi: 10.1002/ctm2.954.

76. Tsushima K, Bugger H, Wende AR, Soto J, Jenson GA, Tor AR, McGlauflin R, Kenny HC, Zhang Y, Souvenir R, Hu XX, Sloan CL, Pereira RO, Lira VA, Spitzer KW, Sharp TL, Shoghi KI, Sparagna GC, Rog-Zielinska EA, Kohl P, Khalimonchuk O, Schaffer JE, Abel ED. Mitochondrial Reactive Oxygen Species in Lipotoxic Hearts Induces Post-Translational Modifications of AKAP121, DRP1 and OPA1 That Promote Mitochondrial Fission. Circ Res 122: 58–73, 2018. doi: 10.1161/CIRCRESAHA.117.311307.

77. Hinton AOJr, Yang Y, Quick AP, Xu P, Reddy CL, Yan X, Reynolds CL, Tong Q, Zhu L, Xu J, Wehrens XHT, Xu Y, Reddy AK. SRC-1 Regulates Blood Pressure and Aortic Stiffness in Female Mice. PLOS ONE 11: e0168644, 2016. doi: 10.1371/journal.pone.0168644.

78. Pathak GA, Singh K, Wendt FR, Fleming TW, Overstreet C, Koller D, Tylee DS, De Angelis F, Mendoza BC, Levey DF, Koenen K, Krystal JH, Pietrzak RH, O’ Donell C, Michael Gaziano J, Falcone G, Stein MB, Gelernter J, Pasaniuc B, Mancuso N, Davis LK, Polimanti R. Genetically regulated multi-omics study for symptom clusters of posttraumatic stress disorder highlights pleiotropy with hematologic and cardio-metabolic traits. Mol Psychiatry 27: 1394–1404, 2022. doi: 10.1038/s41380-022-01488-9.

