## Supplementary figures and images for "Three-Dimensional Mitochondria Reconstructions of Murine Cardiac Muscle Changes in Size Across Aging"

### Supplemental Figure

| | 3-month (n=9) | 1-year (n=10) | 2-year (n=10) |
|---|---|---|---|
| HR (bpm) | 704.1±38.24 | 697.8±32.92 | 667.5±33.46 |
| EF (%) | 0.8577±0.030 | 0.8435±0.0285 | 0.8454±0.040 |
| CO (µL/min) | 12097±2729 | 19502±4959 | 24867±9793 |
| SV (µL) | 17.28±4.15 | 27.92±6.720 | 37.25±14.51 |
| Volume Mass | 0.3508±0.06821 | 0.3693±0.06881 | 0.3741±0.06789 |
| LVT (mm) | 0.717±0.05 | 0.826±0.055 | 0.998±0.172 |
| EAS (mm) | 0.68±0.10 | 1.01±0.129 | 1.235±0.50 |
| EAD (mm) | 4.05±0.7270 | 5.63±1.053 | 7.07±2.35 |
| EMS (mm) | 4.87±0.41 | 6.00±0.4380 | 6.40±0.65 |
| EMD (mm) | 5.91±0.4623 | 7.01±0.3983 | 7.36±0.50 |
| ESV | 2.77±0.5 | 5.06±0.908 | 6.76±3.62 |
| EDV | 20.05±4.374 | 32.97±7.183 | 44.01±17.54 |

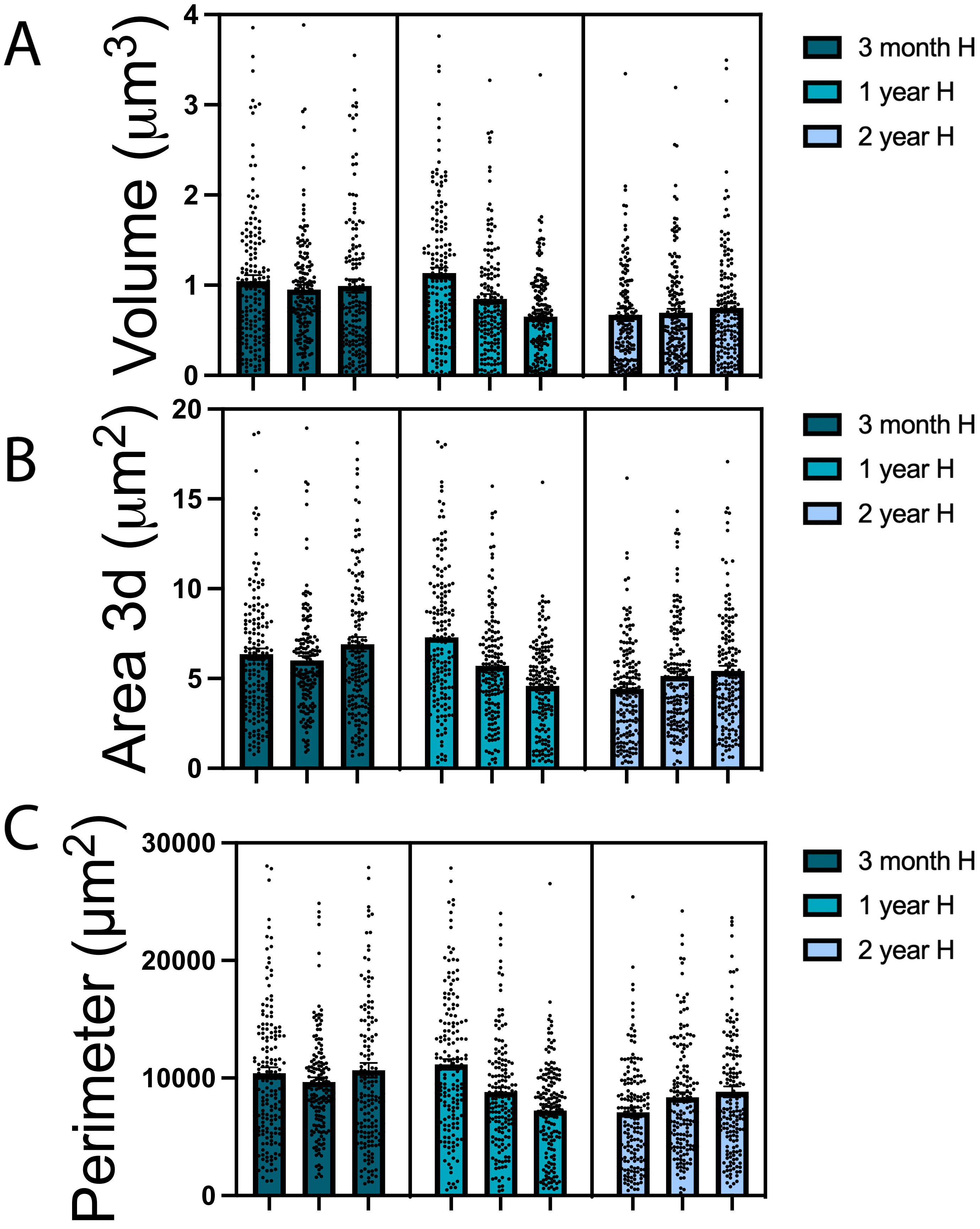

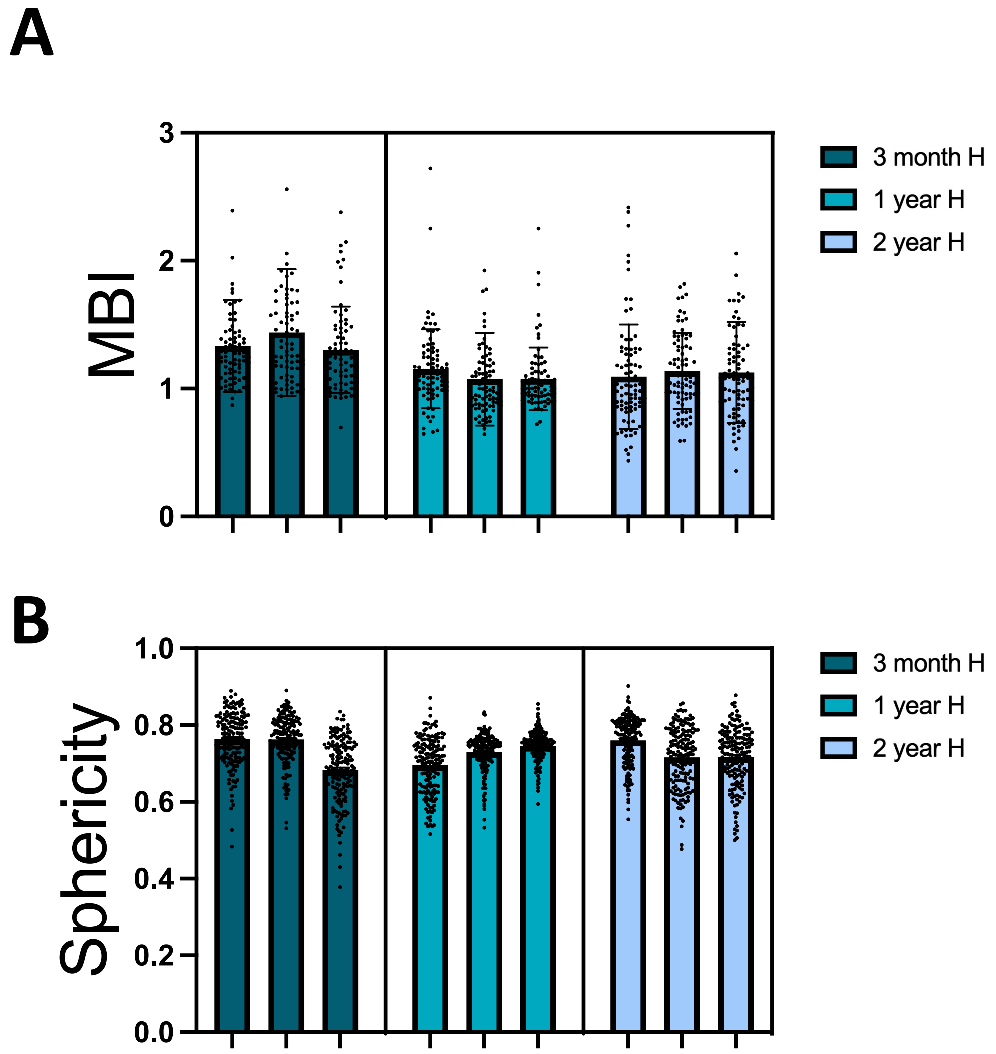

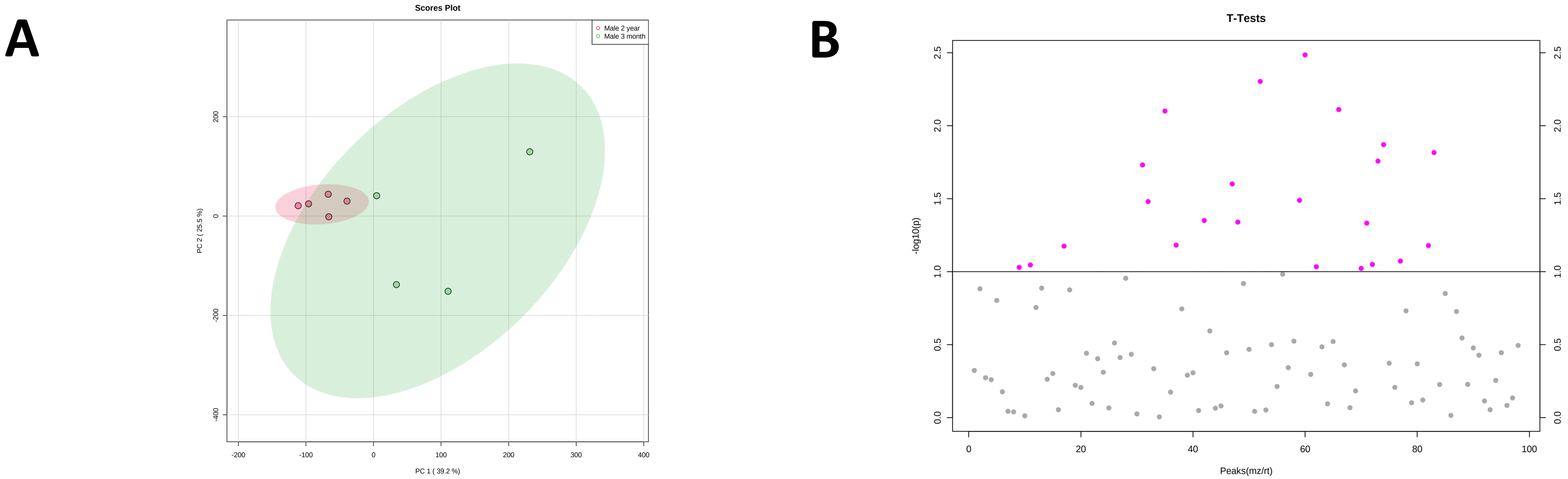
